## Additional results, supplementary figures and tables for "Population genomics and epigenomics provide insights into the evolution of facultative asexuality in plants"

#: these authors contributed equally

*: Corresponding author

### **1. Additional Results**

#### **1.1 Update of *S. polyrhiza* annotation**

Using recently published transcriptomic data and proteomes, we updated the genome annotation of *S. polyrhiza* using Maker (version 2.31.10)^1^ and Braker (version 2.1.5)^2^ (Supplementary Fig. 16). We predicted gene models using isoform sequencing (Iso-seq) reads, short-read transcriptomes from different tissues of *S. polyrhiza* and high-quality proteomes from closely related lineages or well-established model species as empirical evidence (see Section 2.1 of the Supplementary Information). Although the total number of annotated genes only slightly increased (926 genes more than Sp7498V2), the new annotation (SpGA2022) reaches higher completeness and is less fragmented than the previous annotation (Supplementary Table 6 and Fig. 17). The DOGMA^3^ score increased from 71.9 to 79.3 (Supplementary Fig. 18).

#### **1.2 MADS-box transcription factor family in *S. polyrhiza***

The MADS-box family controls different developmental processes, including root, leaf, flower, and fruit development^4,5^. *Spirodela polyrhiza* has lost many MADS-box TFs clades^6^, which might be associated with the reduced size and simplified architecture^7^. Here, we reannotated 43 MADS-box genes in the *S. polyrhiza* genome (Supplementary Fig. 19). Based on the phylogenetic relationship, we classified these MADS-box TFs into 14 clades (Supplementary Table 7), three from the Type I subfamily (Supplementary Fig. 20), and 11 from the Type II subfamily (Supplementary Fig. 21). Consist with the previous study^7^, we found that *AGL15*-, *FLC*-, *AGL9*- *AGL12*-, and *OsMADS32*-like clades are missing in the *S. polyrhiza* genome.

#### **1.3 Structure variation in the *S. polyrhiza* population**

We characterized genome-wide structural variations (SVs) using an adopted joint genotyping pipeline^8^ (Supplementary Fig. 22). Using long reads from genotype SP014 (strain 9509)^9^, we evaluated the performed precision-recall (PR) of our pipeline. The precisions of insertions and deletions identified based on short reads reached 28.0% and 64.5%, respectively, which are comparable with that found in a recent study focused on soybean population genomics^10^. The high consistency of population structures estimated based on SVs and SNPs often indicates high accuracy of SV characterizations^11,12^. In the PCA analysis using the identified SVs, we found that four populations were separated according to the first three PCs (Supplementary Fig. 23), consistent with the pattern observed using SNPs. Using fastStructure with SVs, we also identified four populations (Supplementary Fig. 24) with the same genotype assignment as with the analysis using SNP data. The clustering pattern using SVs and SNPs also showed similar patterns (Supplementary Fig. 25).

In total, we identified 2,089 deletions, 825 insertions, and 291 duplications in the *S. polyrhiza* genomes. The average size of SVs in *S. polyrhiza* is 0.0024/100 bp, which is significantly less than in other studied plants^11,12^.

#### **1.4 Population demographic history**

We developed three competing demographic models using coalescent simulations to provide further insights into the population’s demographic history.

*Model choice*

Our ABC-based model selection procedure favours Model 3 (Posterior Probability; P=0.88) as the most probable model, compared to Model 1 (America-origin, P=0.027) and Model 2 (Asia-origin, P=0.097) (Supplementary Fig. 3). this result implies that the sampled American and SE-Asian populations are equally distant from a putative ancestral population, and this is consistent with the phylogenetic observation, where the branch lengths of the American and the SE-Asian clades are very similar.

*Parameter inference*

After performing ABC-based parameter inference on Model 3, we found that the SE-Asian population has the largest *Ne*, followed by the American, Indian, and European populations (Supplementary Table 9). Concerning the colonization times, both our sampled American and SE-Asian populations split from an ancestral population ca. one million generations ago. Then, the Indian and European populations split from the SE-Asian branch ca. 51,000 and 12,000 generations ago, respectively (Supplementary Table 9). Finally, migration in and out of Europe is the highest, followed by migration between Asia and America and in and out of India (Supplementary Table 9).

#### **1.5 Population/branch-specific selection scans**

Selection scans with 3P-CLR (using the American population as an outgroup) identified a total of 1883 genes on the SE-Asian branch, 593 genes on the Indian branch, and 401 genes on the European branch (Fig. 4a), corresponding to the top 1% of CLR values on each population branch (Supplementary data 5). Of these genes, 77 were common between Europe and India, 18 between SE-Asia and Europe, 32 between SE-Asia and India, and 12 in all three populations (Fig. 2b and Supplementary data 5). We tested the significance of the largest gene overlap (India and Europe) in comparison with the genomic background (all annotated genes from 20 chromosomes) and found a *P-value* < 2.2e-16 (Fisher’s Exact Test). In addition to the three genes mentioned in the main text, [SpGA2022_013448 (*FLK*-like), SpGA2022_006111 (*BB*-like) and SpGA2022_055195 (*CYP78A9*-like)], several other genes are also likely involved in flower and organ development. SpGA2022_006114 (chromosome 3), annotated as *PFP*, is a phosphofructokinase with higher expression in flowers and fruits than in roots and leaves^13^. SpGA2022_052120 (chromosome 4), annotated as *ARF1* Binding protein, encoding an auxin response factor, and is involved in floral organ abscission and positive regulation of flower development^14^. SpGA2022_016777 (chromosome 13), annotated as *SOMBRERO*, is involved in auxin signalling, negatively affecting the cellular re-specification at the root tip, where *SOMBRERO* orchestrates both the formation of extra root cap layers and primary root growth under phosphate scarcity^15^.

In addition to the five MADS-box genes that were under positive selection in the Indian population, two MADS-box genes (SpGA2022_013026 and SpGA2022_013226) were also under selection in the SE-Asian population.

### **2. Additional Methods**

#### **2.1 Genome annotation update**

To improve the *S. polyrhiza* genome annotation, we developed a pipeline integrating short- and long-read transcriptomic data, available proteomes, and *ab initial* gene predictions (Supplementary Fig. 17). RNA-seq short reads from *S. polyrhiza* frond, root, and whole plant^16^ were downloaded from NCBI and filtered using Skewer (v0.2.2)^17^. After the filtration, they were mapped to the reference genome of *S. polyrhiza* using HISAT2 (v2.2.1)^18^ with the “--dta” mode. SAMtools (v1.10)^19^ was used for bam file sorting and indexing. Scallop (v0.10.5)^20^ was used to assemble the transcripts. Iso-seq data^16^ were downloaded from NCBI, and the full-length transcripts were aligned to the *S. polyrhiza* reference genome using minimap2 (v2.21)^21^ under the “-x splice:hq” mode. Only mapped transcripts were kept as empirical evidence in the Maker pipeline.

High-quality proteomes from *Oryza sativa* (v7.0, Phytozome 13)^22^, Maize (AGPv3.22, MaizeGDB)^23^, *Arabidopsis thaliana* (TAIL10)^24^, and *Zostera marina* (V2, OrcAE)^25^ were included as the empirical evidence for the Maker pipeline to predict gene models.

We combined Maker (version 2.31.10)^1^ and Braker (version 2.1.5)^2^ pipelines for annotating the gene models (Supplementary Fig. 17). One round of SNAP *ab initio* gene prediction was performed based on the repeat soft-masked reference genome of *S. polyrhiza*. Results from the Braker pipeline and the previous annotation Sp7498V2 were provided to Maker as “legacy annotations”. After that, the transcript model from the iso-seq data was used to correct the gene models. BLAST+ (v2.11.0)^26^ and InterProScan (version 5.50-84.0^27^, running under Java/11.0.2) were used to perform functional and Gene Ontology (GO) annotation (Supplementary Fig. 17).

**2.2 Identification of the MADS-box transcription factors**

All protein-coding genes from SpGA2022 annotation were aligned with the MADS-box genes from *O. sativa*^28^ and *A. thaliana*^24^ using BLAST+ (v2.11.0)^26^ with an e-value threshold of 1e-5. We search against the SRF- and MEF2-Type MADS domains using “hmmsearch” from HMMER (v3.3.2, http://hmmer.org/)^29^. Results from those two processes were combined and further filtered using the NCBI Conservative Domain Database (CDD)^30^ (Supplementary Fig. 20). As a result, 43 MADS-box genes were identified in *S. polyrhiza* (Supplementary Data 6).

To annotate the identified 43 *S. polyrhiza* MADS-box genes, we collected annotation information based on either TAIR (www.arabidopsis.org) or previous studies of the MADS-box gene family in *A. thaliana*^31^, *O. sativa*^32^, *Ananas comosus*^33^, *Saccharum spontaneum*^34^, and *Nelumbo nucifera*^35^. In total, 223 MADS-box gene coding proteins from *A. thaliana*, *O. sativa*, and *S. polyrhiza* were aligned using MAFFT (v7.490)^36^. Phylogenetic trees of type I and type II MADS-box families were reconstructed using FastTree (v2.1.11)^37^ with default parameters. According to the phylogenetic clustering, 43 MADS-box genes were then classified into 14 clades (Supplementary Table 7). Tree visualization and annotation were done using the “ggtree” package (v3.2.1)^38^ with R (v4.1.0).

**2.3 Structure variations identification in *S. polyrhiza***

We adopted the joint genotyping pipeline (Supplementary Fig. 23) from Eggertsson *et al.*^8^ to call SVs. Compared to the original method in which Manta was the only SV caller, we added another four popular SV callers in the pipeline. We found that this method could make full use of Manta’s high sensitivity while keeping the false positives relatively low. Further validation based on the long-read sequencing data suggested that this method provided the most confident SV dataset when compared with other prevailing methods.

For each *S. polyrhiza* sample, Manta^39^, Smoove (Lumpy calling and svtyper genotyping, <https://github.com/brentp/smoove>)^40,41^, GRIDSS^42^, Delly^43^, and SvABA^44^ were used independently to call individual-level SVs. After that, all SVs from each caller were merged using svimmer’s “join mode”. For each sample, we kept only SVs called by both Manta and at least one other caller. After that, we generated a population-level SV call set using svimmer, which was then used for genotyping with GraphTyper2^8^.

We applied several filtrations to the population-level genotyped SV set. First, complex SVs and SVs from organelle genomes were removed. Second, SVs with a size less than 50 bp or larger than 500 kb were removed, as small SVs were included in the GATK pipeline, and large SVs were difficult to confirm. Third, deletions that contain assembly gaps were removed due to uncertainties in the genome quality; Fourth, SVs with minor allele frequency lower than 0.01 or higher than 60% of missing genotypes were removed; Fifth, for duplications, only genotypes with “Genotyping Quality (GQ)” higher or equal to 20 were kept. We didn’t apply the default genotyping filtration from GraphTyper2 because its strict filtration criteria would remove a high proportion of positive calls^8^. We used BCFtools^45^, bgzip, and tabix^46^ for most of the Variant Call Format (VCF) file manipulations. The filtrations were done using BCFtools, VCFtools, and VcfFilter (<https://github.com/biopet/vcffilter>).

To estimate population structure using SVs, we converted the VCF file into PLINK format using VCFtools^47^. The PCA was carried out using PLINK, while the population structure analysis was estimated using fastStructure. For the SV-based phylogenetic tree reconstruction, we converted SVs into a p-distance matrix using VCF2Dis (<https://github.com/BGI-shenzhen/VCF2Dis>). The PHYLPNEW program from EMBOSS^48^ was used to build the neighbour-joining tree. The R package “ggtree” (v3.2.1)^38^ was used for tree visualizations. SV alignments were visualized using Samplot^49^, and gene models were plotted using JBrowser2^50^.

**2.4 Demographic analysis**

To infer the demographic history of our sampled populations of *S. polyrhiza,* we used Approximate Bayesian Computation (ABC), which makes use of observed data, candidate demographic models, and simulations of the observed data under the candidate models to infer population demographic parameters such as population sizes, migration rates, and colonization times.

Data consists of SNPs coming from intergenic regions of the four populations (SE-Asia, North America, Europe, and India) of *S. polyrhiza*. From each population, we selected 17 non-clonal individuals (the minimum number observed, which corresponds to the American population) to ensure all populations have the same number of individuals for downstream analyses. To select intergenic regions for demographic inference, we performed the following steps. First, we obtained the coordinates of all intergenic loci based on our latest annotation. Then, we removed all loci that overlapped with structural variants. Then, we sorted the loci by size and by chromosome and selected the three largest loci from each chromosome. We manually inspected the resulting alignment for each locus and discarded the loci with a large percentage of missing data or no SNPs available. A total of 39 intergenic loci passed all the filters described above, averaging two loci per chromosome (Supplementary Table 10). This filtering procedure is ideal for demographic inference because it keeps a few loci per chromosome (thus increasing the independence between loci, which is required in coalescent simulations) while keeping a large number of SNPs. All 39 intergenic loci yielded a total of 13,065 SNPs for demographic inference. From each of these loci, we calculated the following summary statistics: number of segregating sites S, Waterson’s theta, pi, Tajima’s D, Fu and Li’s D, Fay and Wu’s H, average LD (ZnS), the site-frequency spectrum (SFS) per population, the joint SFS (JSFS) for each pair of populations, and the summaries of the JSFS as described by Wakeley and Hey^51^.

We tested three demographic models (Supplementary Fig. 3). Model 1 consists of an American origin, with subsequent colonization of Asia, and then from Asia independent colonization of Europe and India. Model 2 starts with Asia as the ancestral population, then colonization to America and further independent colonization of Europe and India from Asia. Model 3 consists of a putative ancestral population, then colonization to America and Asia, and subsequent colonization of India and Europe from Asia. Migration was allowed between all current populations.

For each of the models described above, we performed 50,000 coalescent simulations with the software *msms*^52^. This program simulates coalescent trees for each population under any given demographic model while accounting for selection. The output of *msms* are SNPs from which we calculated the following summary statistics: number of segregating sites S, Waterson’s theta, pi, Tajima’s D, Fu and Li’s D, Fay and Wu’s H, average LD (ZnS), the site-frequency spectrum (SFS) per population, the joint SFS (JSFS) for each pair of populations, and the summaries of the JSFS as described by Wakeley and Hey^51^. These same statistics were calculated for the observed loci described below.

The model choice was performed within an ABC framework. Posterior probabilities for each model were calculated according to Fagundes *et al.*^53^. Model choice was done based on the mean and variance (across loci) of the number of segregating sites *S*, Tajima’s *D*, linkage disequilibrium (*Z*_nS_), and population differentiation statistics in all four populations. In our analysis, Watterson’s Θ_W_, Π*_n_*, and *K* were correlated with *S_n_*, and therefore its inclusion does not change the results of the model choice procedure. When comparing all models, the model with the highest posterior probability was chosen as the best fit for the observed data.

On the best model, the inference was based on ABC rejection^54,55^ and regression^56^ methods. Both methods were performed using *ABCtoolbox*^57^ and checked with Csilléry’s abcR^58^. First, we pooled all statistics and checked for correlations with the parameters. We did not keep statistics that did not correlate with any parameter because keeping them does not provide information for the estimation and would only add noise to the final estimates. All these statistics were transformed using partial least squares (p.l.s.) as implemented in Wegmann *et al.*^57^. This transformation is advantageous because it extracts a small number of orthogonal components from a higher dimensional array of summary statistics. The new set of transformed statistics (with reduced dimensionality) reduces the noise produced by uninformative summary statistics. Moreover, the p.l.s.-transformed statistics are completely uncorrelated with one another, ensuring the assumption of singularity, which is required for estimating parameters according to the regression method^56^.

**3. Supplementary Figures**

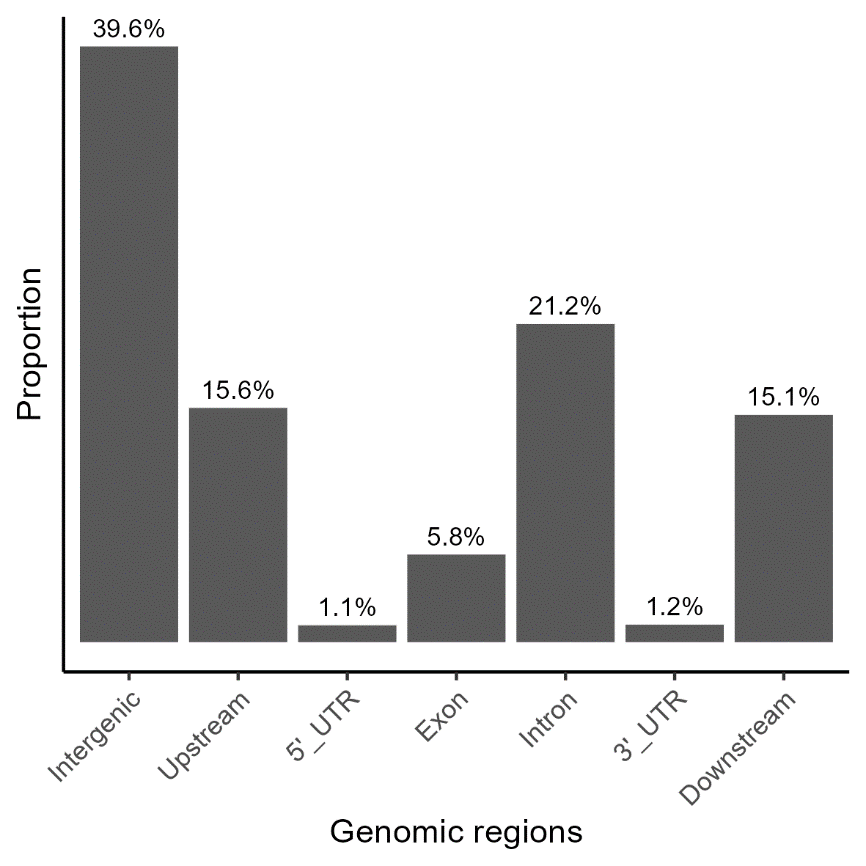

Supplementary Fig. 1. SNP distributions among different genomic regions.

The annotation of SNPs was calculated using SnpEff.  “Upstream” and “downstream” indicate the flanking 2 kb regions of annotated genes. The intergenic region did not take upstream and downstream information into account.

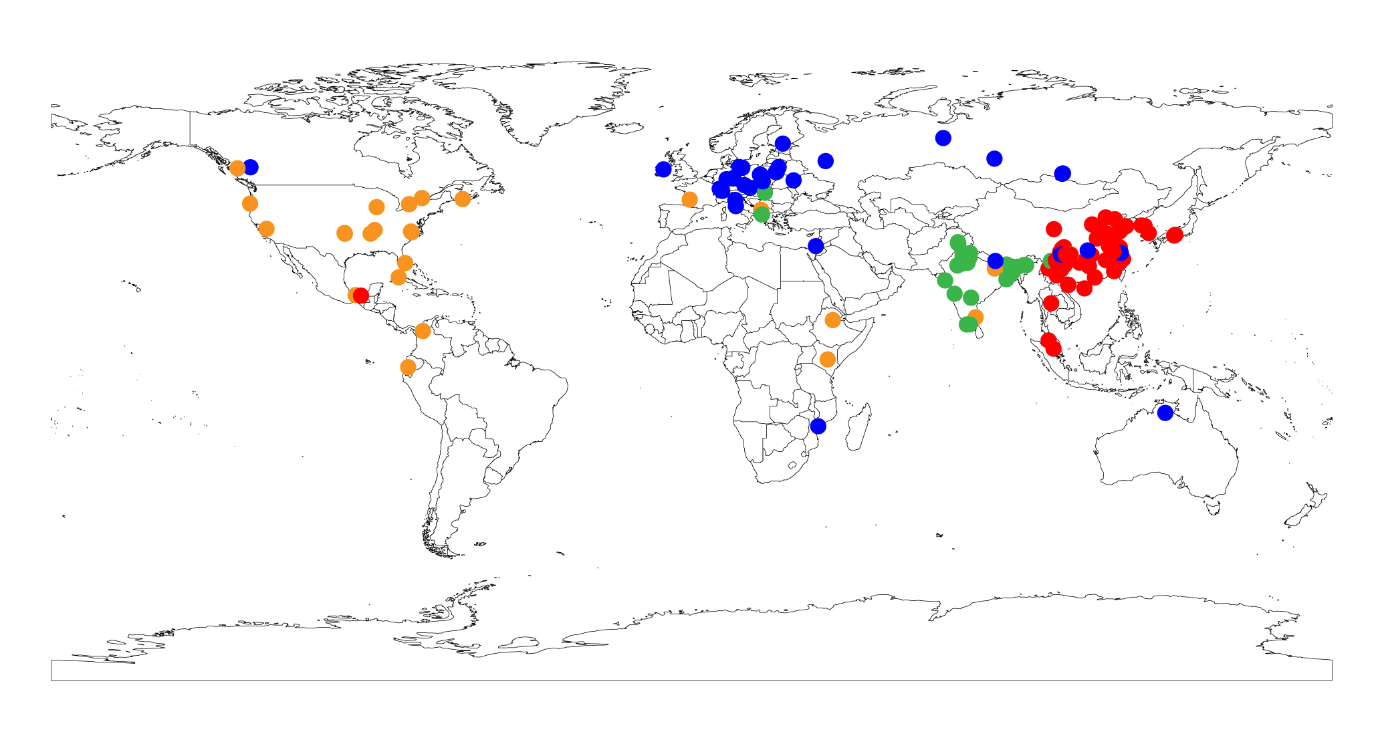

Supplementary Fig. 2. Worldwide geographic distribution of 228 *S. polyrhiza*.

Circular dots indicate the geographic locations from where the samples were collected. Those dots are coloured according to the genetic population stratification: SE-Asian population (red), Indian population (green), European population (blue), and American population (yellow).

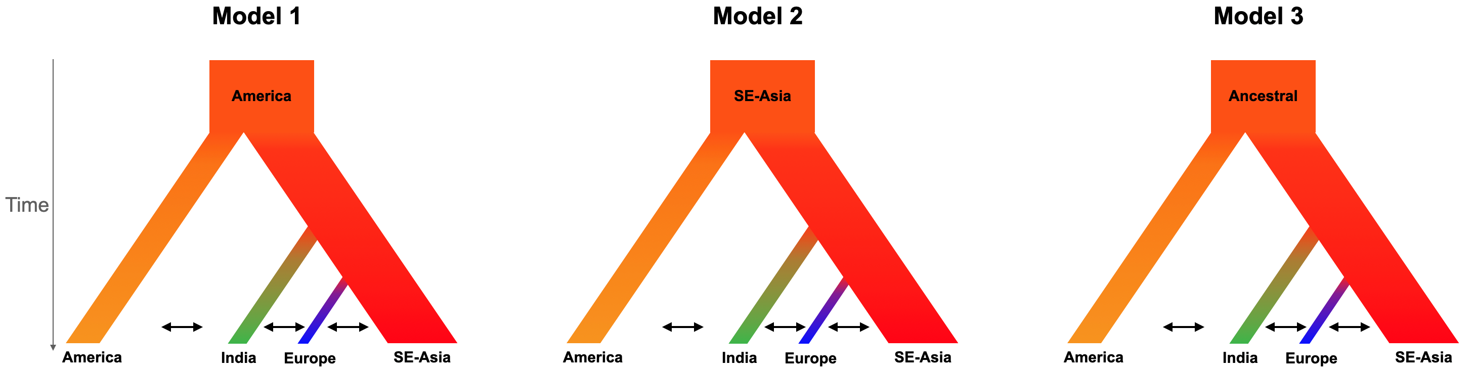

Supplementary Fig. 3. Demographic models tested with ABC.

Model 1 (American origin) had a posterior probability of P=0.027, Model 2 (SE-Asia origin) P=0.097, and Model 3 P=0.88.

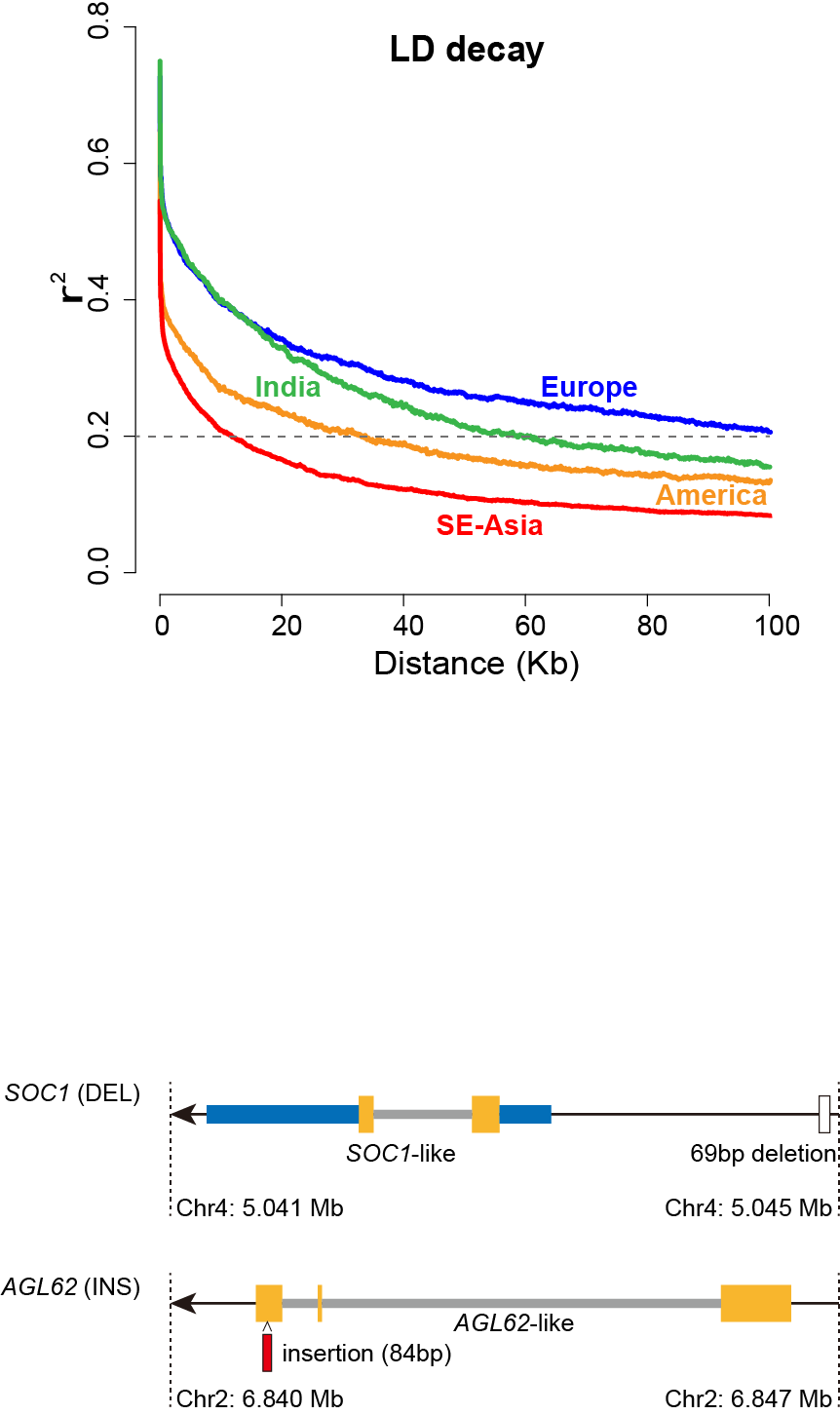

Supplementary Fig. 4. The decay of linkage disequilibrium in *S. polyrhiza*.

Curved lines in different colours indicate the LD decay in all four populations. The grey dashed line represents an r^2^=0.2 used for comparison.

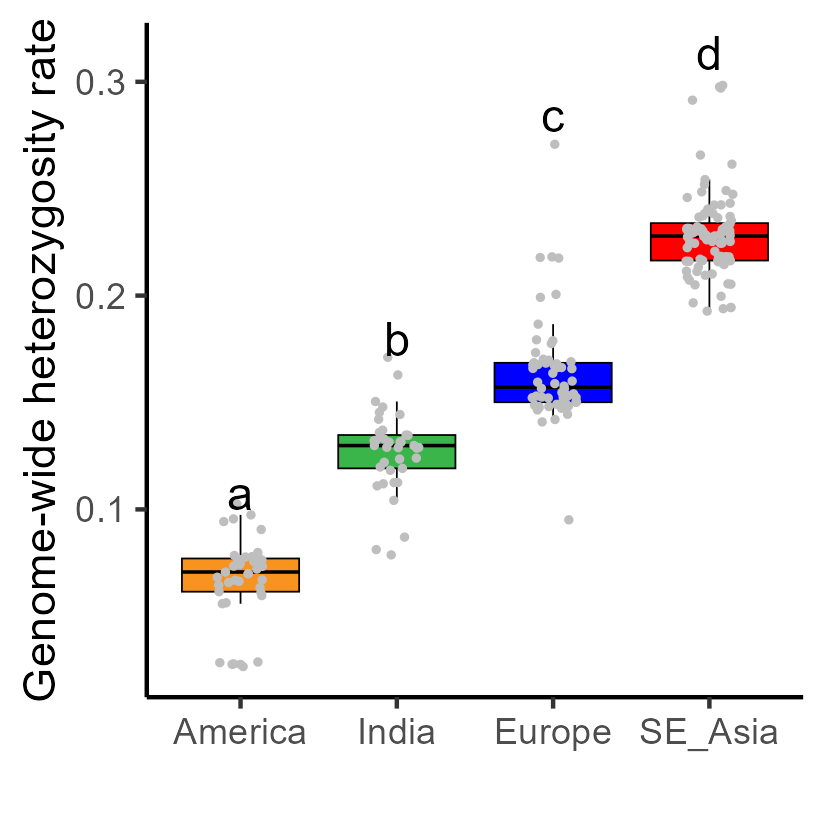

Supplementary Fig. 5. Comparison of genome-wide heterozygosity rate among four populations.

Grey dots indicate the intra-individual heterozygosity rate of each genotype. Lowercase characters show the significance tested using the Wilcoxon test.

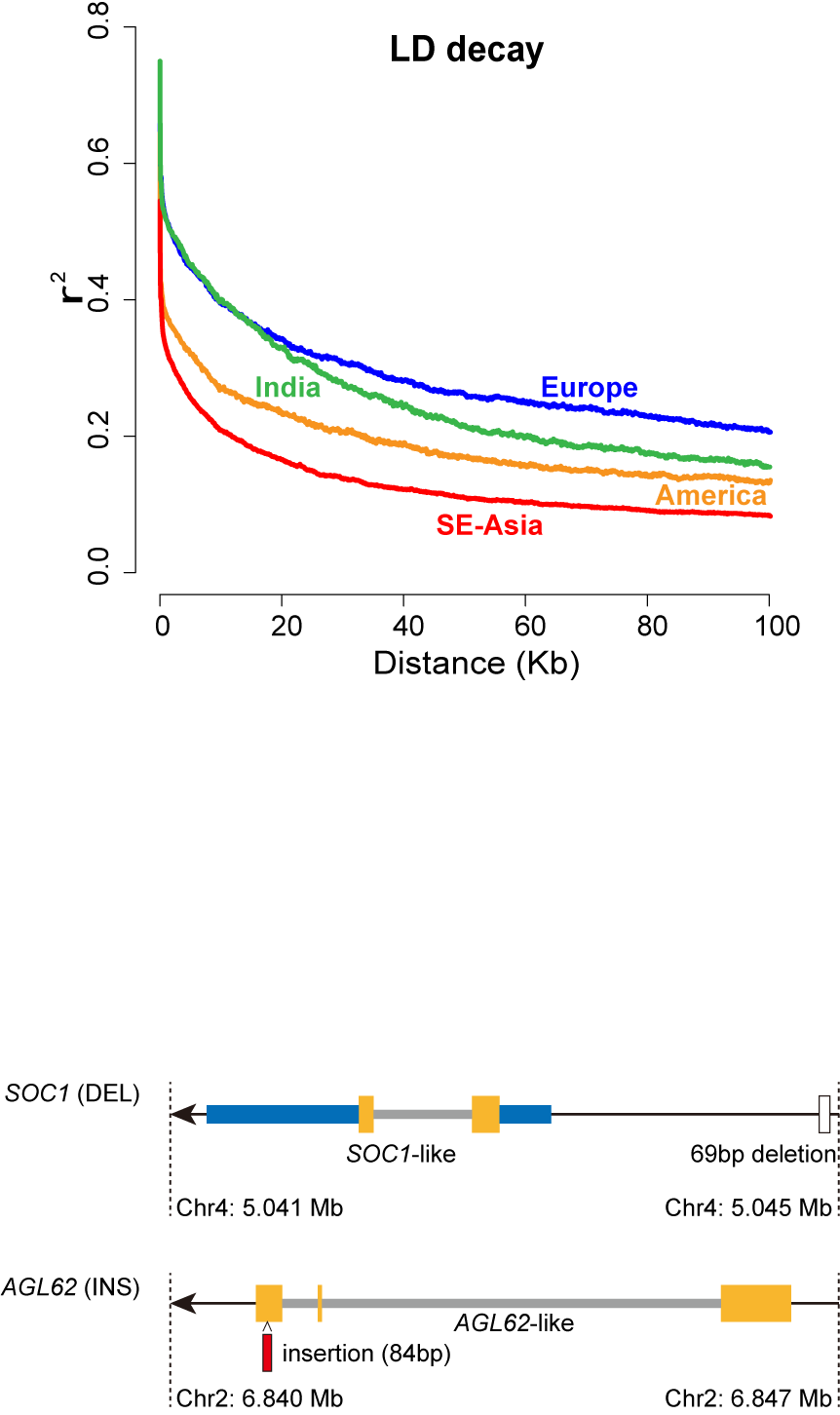

Supplementary Fig. 6. Schematic of the SVs impacting on two MADS-box genes.

SVs affect two MADS-box genes, SpGA2022_007306 (*SOC1-like*) and SpGA2022_005278 (*AGL62-like*), that are related to sexual reproduction. The yellow blocks indicate the exons, the blue blocks indicate the UTRs, and the grey ones indicate the introns. The white rectangle indicates the location of the deletion variation, while the red one indicates the insertion.

 
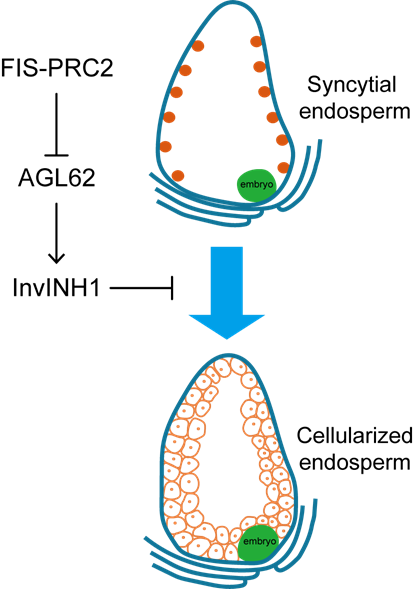

Supplementary Fig. 7. Schematic illustration of the putative *AGL62* pathway.

A black arrow indicates activation, while a bar at the end of a line indicates repression. The thick blue arrow indicates the transition from the syncytial phase to the cellularization phase of the endosperm.

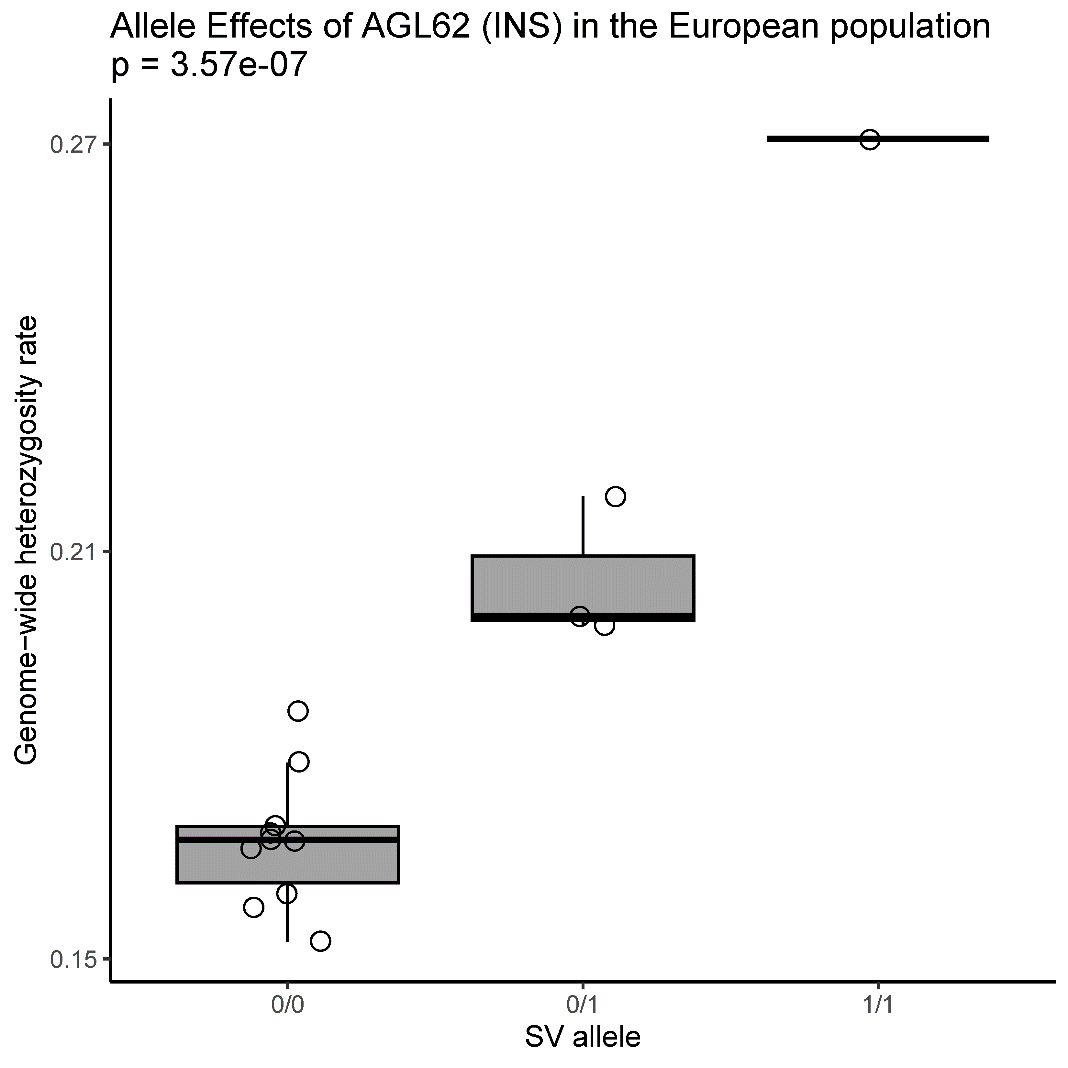

Supplementary Fig. 8. Allele effects of the insertion variation that affect gene *AGL62* in the European population.

Grey circles indicate the intra-individual heterozygosity rate. “0/0” refers to the homozygous reference allele, “0/1” refers to the heterozygous allele, and “1/1” refers to the homozygous alternative allele (allele that is impacted by the SV). P value was estimated from the genetic association using RVTESTS.

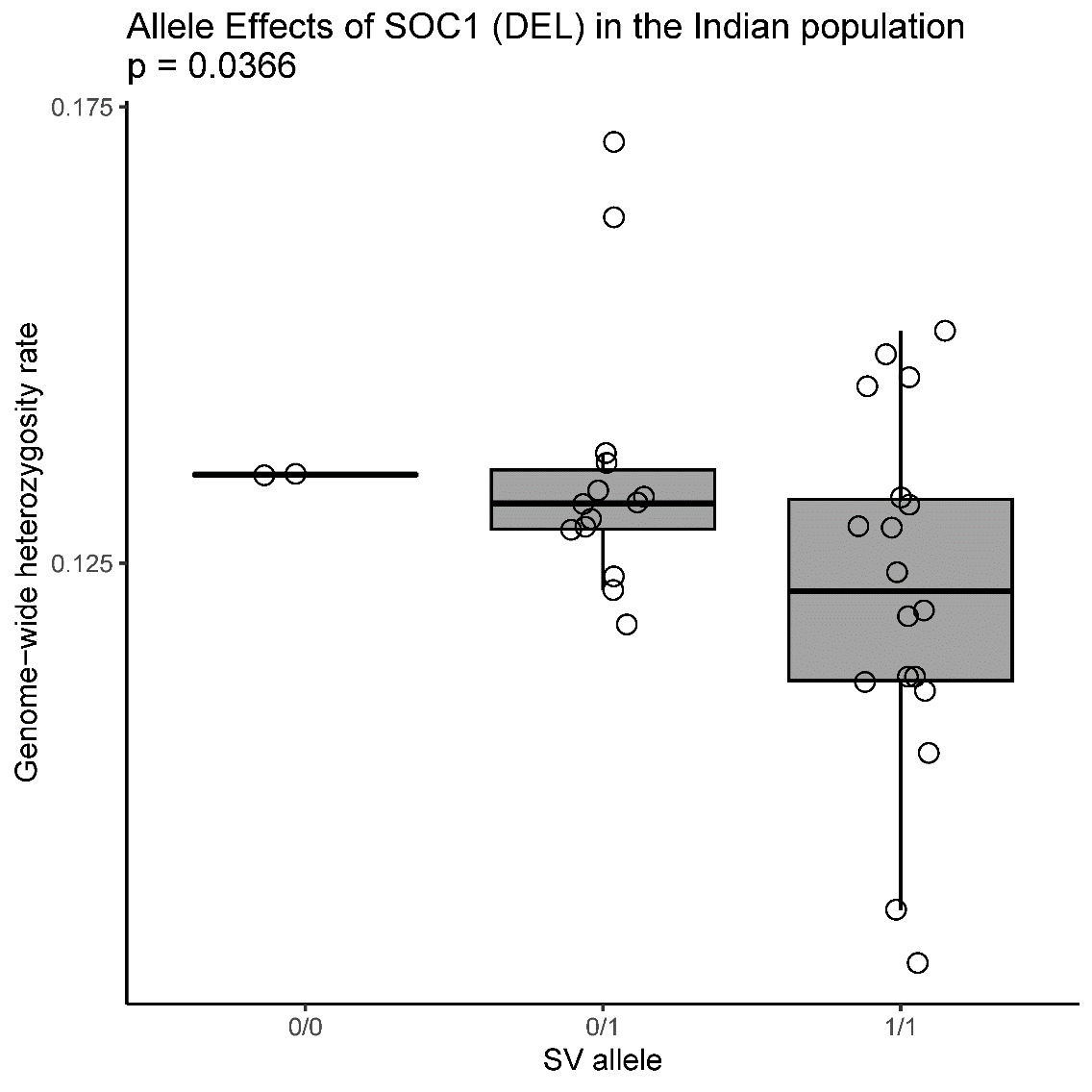

Supplementary Fig. 9. Allele effects of the deletion variation that affect gene *SOC1* in the Indian population.

Grey circles indicate the intra-individual heterozygosity rate. “0/0” refers to the homozygous reference allele, “0/1” refers to the heterozygous allele, and “1/1” refers to the homozygous alternative allele (allele that is impacted by the SV). P value was estimated from the genetic association using RVTESTS.

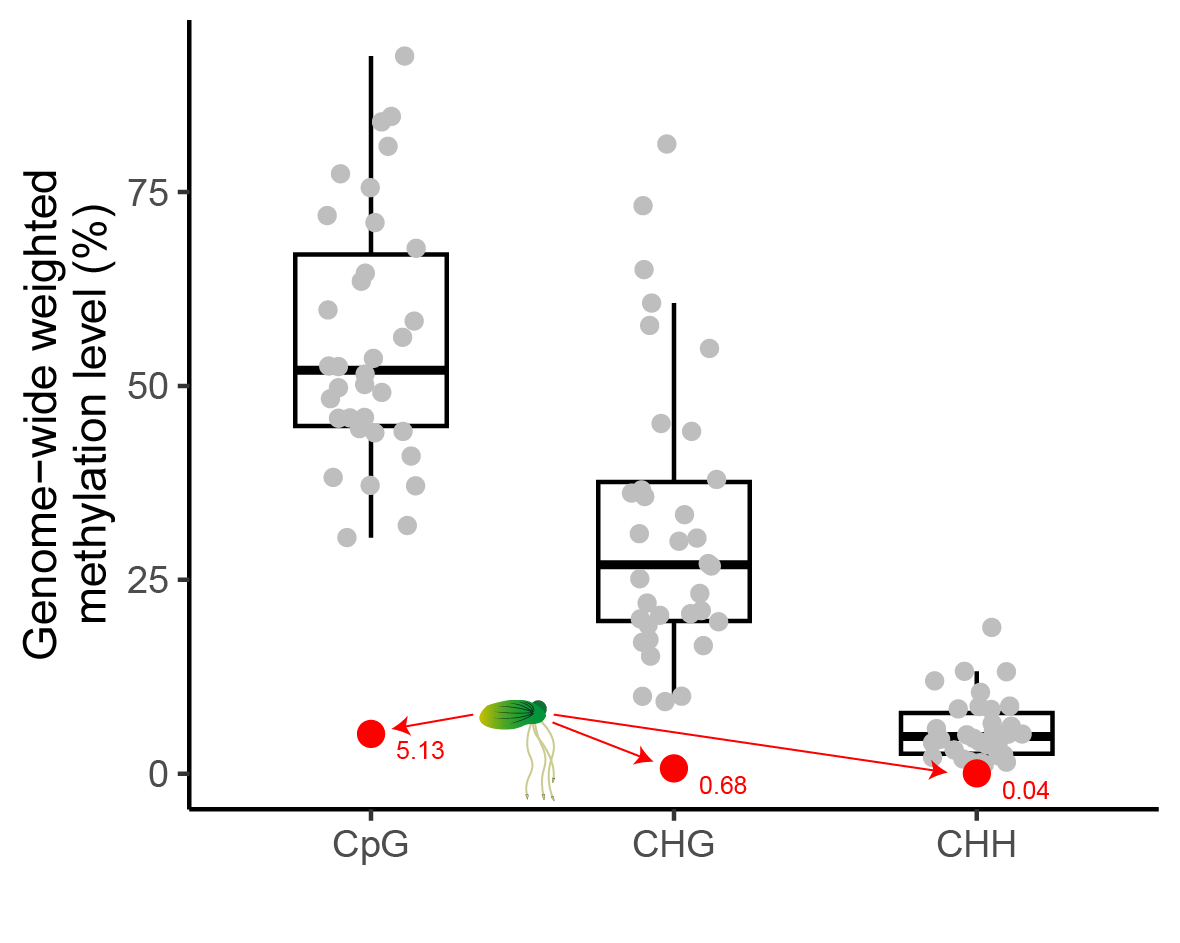

Supplementary Fig. 10. Comparison of genome-wide methylation levels.

The box plot compares the weighted methylation levels of 34 angiosperms and *S. polyrhiza* in CpG, CHG, and CHH. The red dots show the genome-wide weighted methylation levels of *S. polyrhiza*.

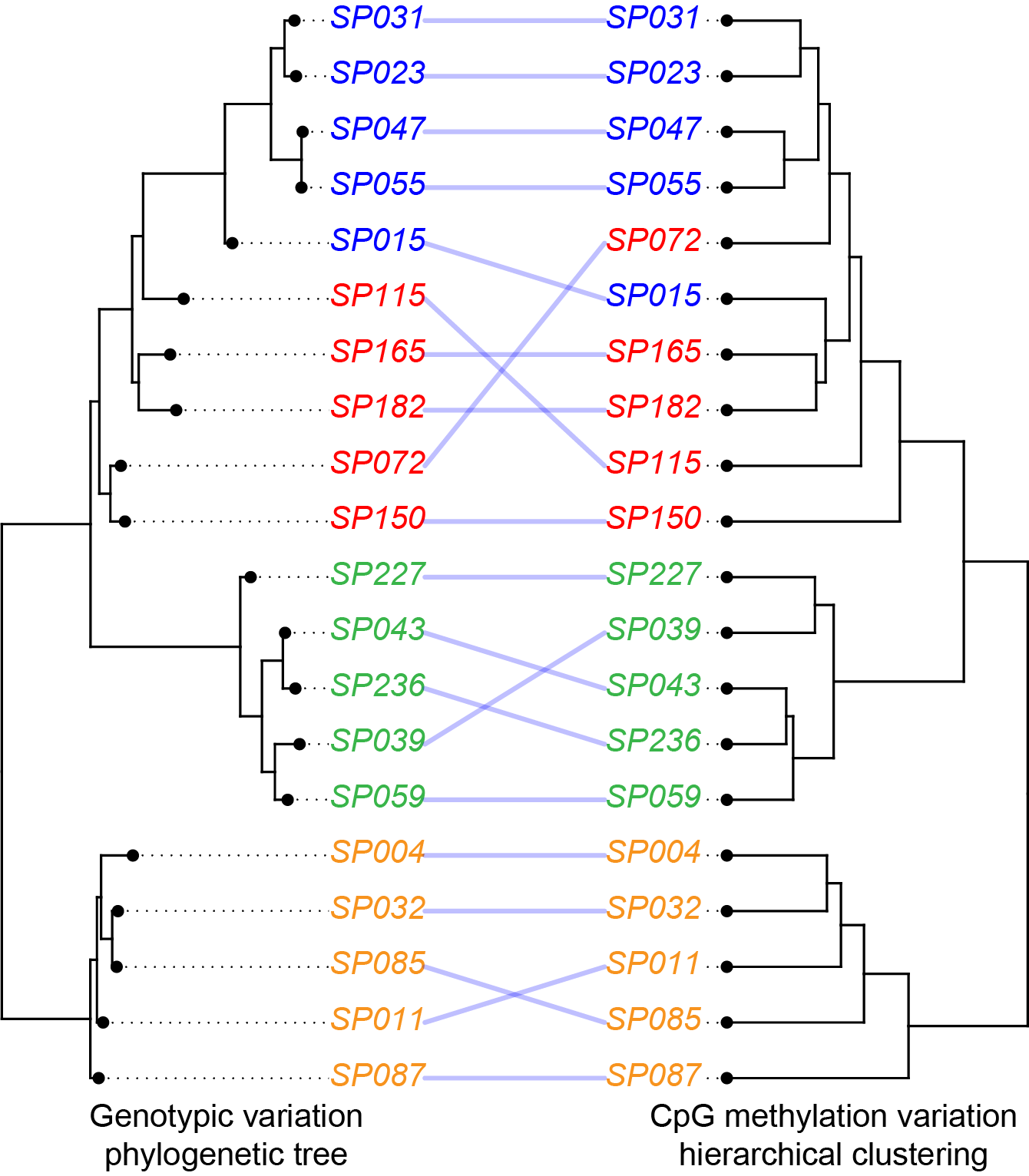

Supplementary Fig. 11. Comparison between the genetic and CpG epigenetic distances among 20 individuals.

The phylogeny was inferred from the SNP data. The hierarchical clustering was calculated based on the genome-wide CpG methylation using “Ward methods”. Tips for both trees were differently coloured based on their population classification.

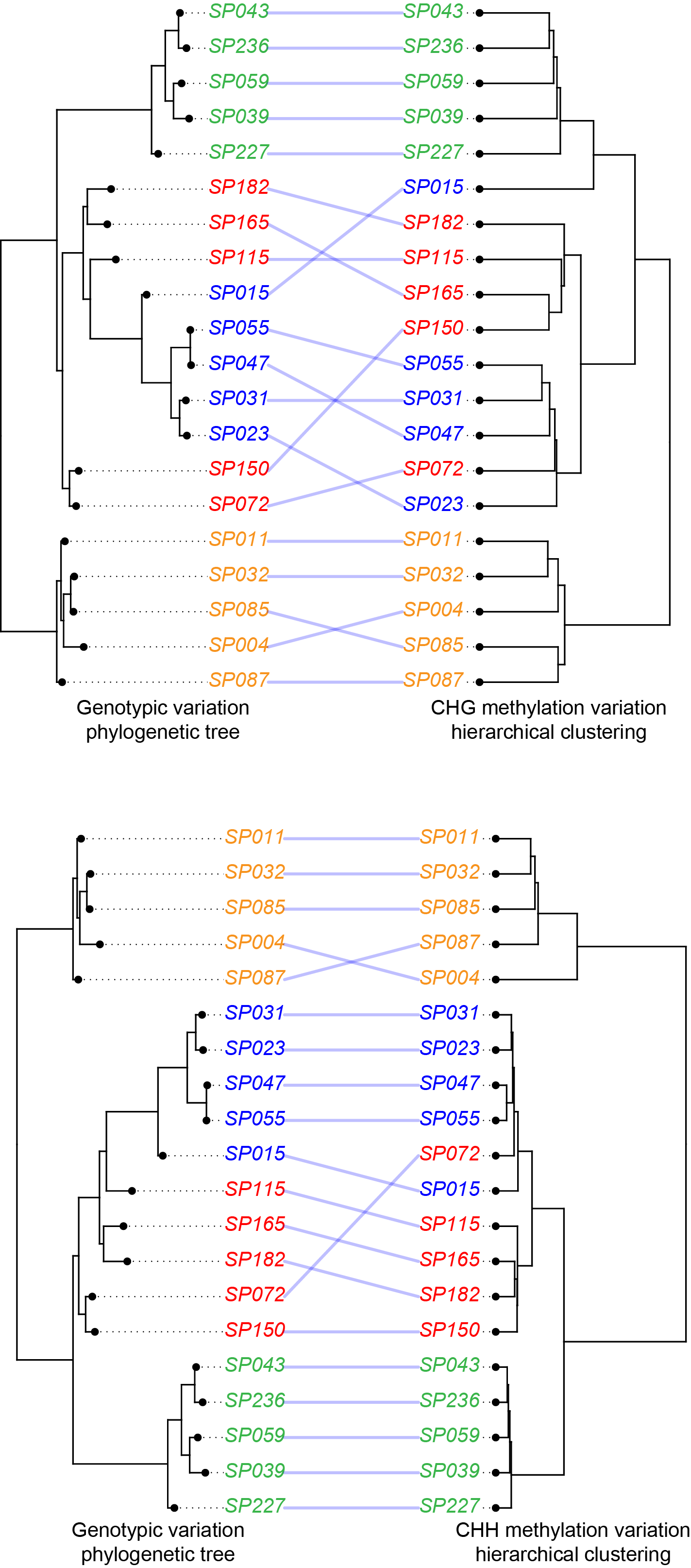

Supplementary Fig. 12. Comparison between the genetic and CHG epigenetic distances among 20 individuals.

The phylogeny was inferred from the SNP data. The hierarchical clustering was calculated based on the genome-wide CHG methylation using “Ward methods”. Tips for both trees were differently coloured based on their population classification.

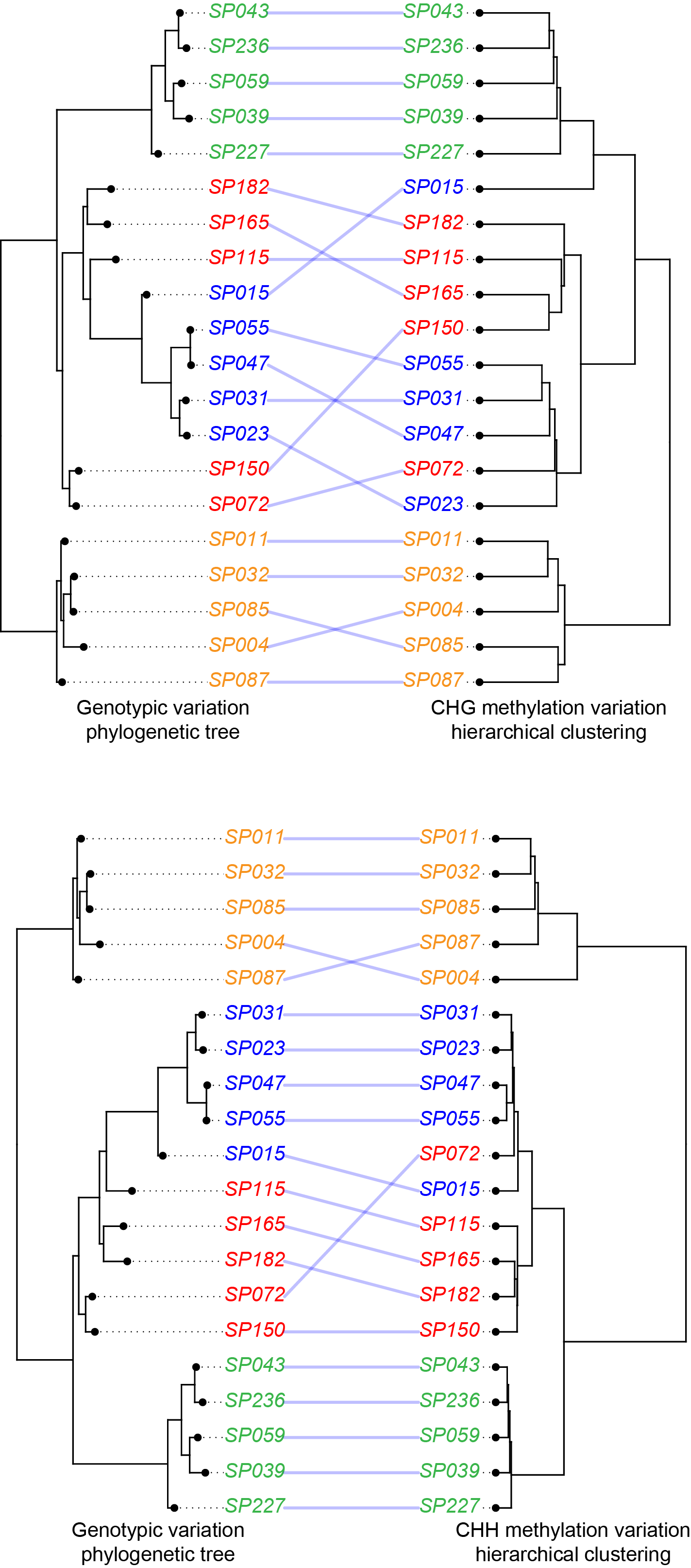

Supplementary Fig. 13. Comparison between the genetic and CHH epigenetic distances among 20 individuals.

The phylogeny was inferred from the SNP data. The hierarchical clustering was calculated based on the genome-wide CHH methylation using “Ward methods”. Tips for both trees were differently coloured based on their population classification.

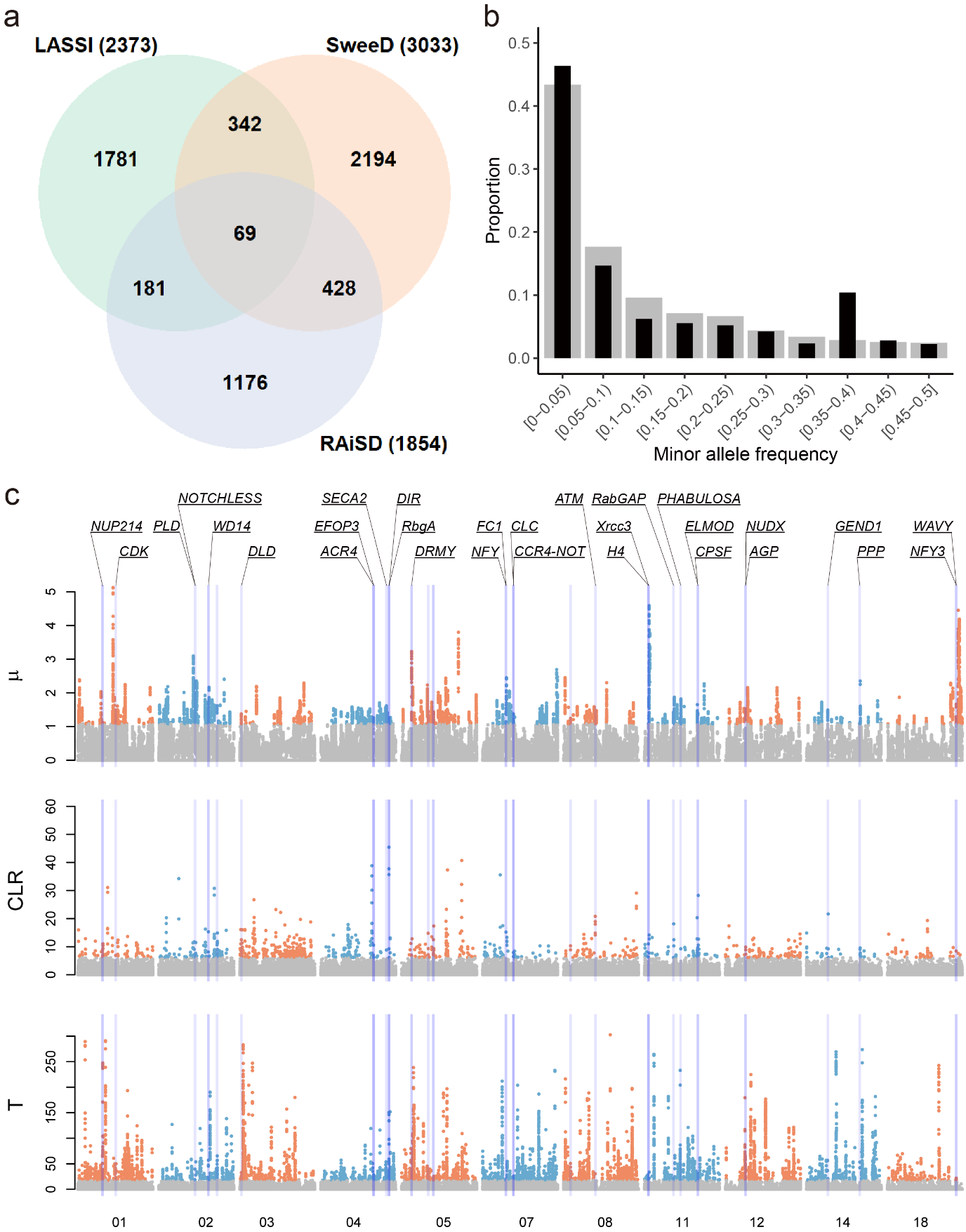

Supplementary Fig. 14. Species-wide selection signature scans.

**a** Number of genes under selection identified using three different methods. In total, 69 genes were found under selection by all three methods. **b** Folded site-frequency spectrum (SFS) calculated from the 69 common genes (black bars) compared to the genome-wide folded SFS (grey bars). **c** Per-chromosome genome-scan results of all three methods, with indications of the genes under selection. Each dot represents a chromosomal locus reported by RAiSD (panel 1), SweeD (panel 2), or LASSI (panel 3) for which the statistic from each software were calculated (µ for RAiSD, CLR for SweeD, and T for LASSI). Significant outliers are shown in colored dots. Purple bars indicate the location of genes which are found to be under selection by all three methods.

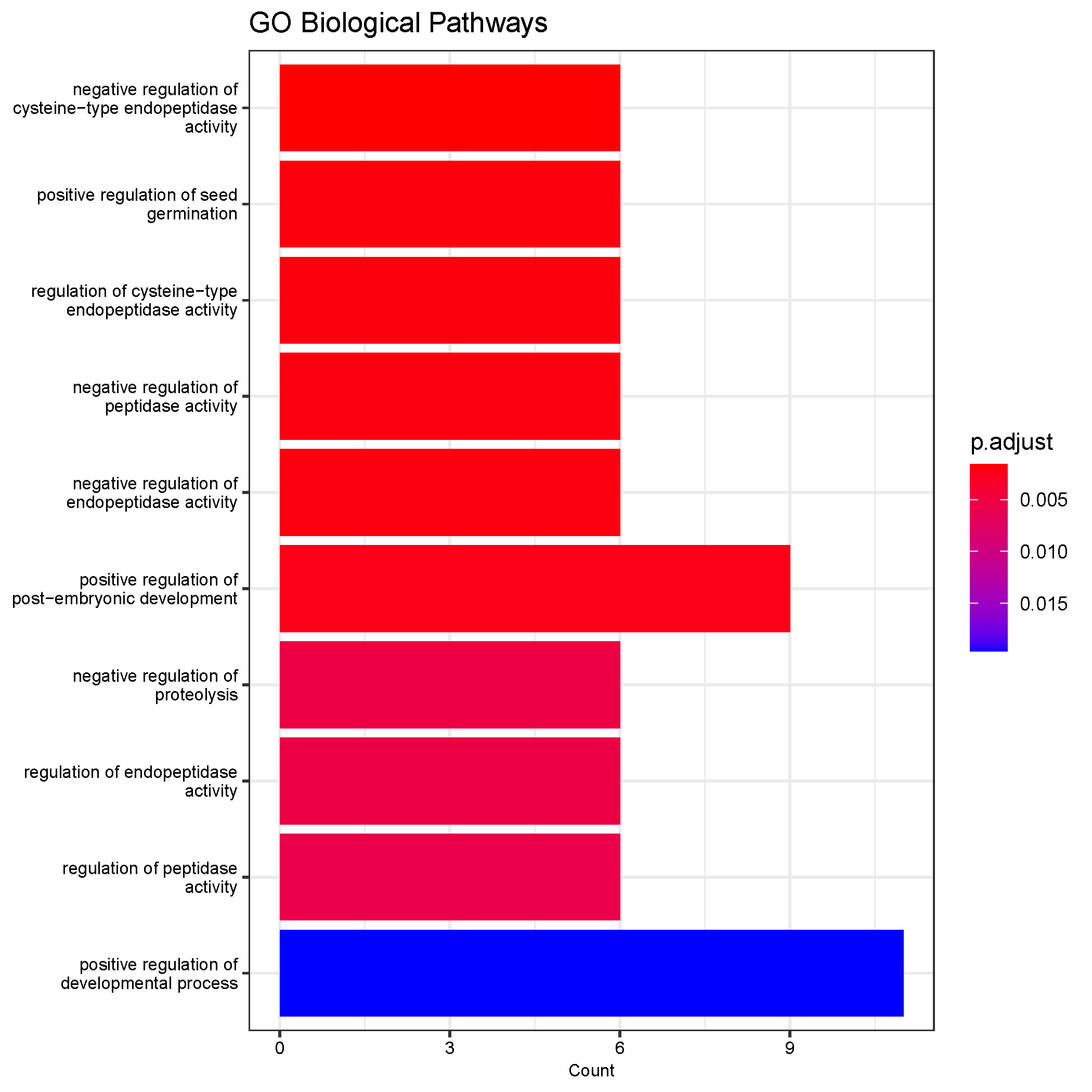

Supplementary Fig. 15. GO enrichment of the genes found by the population-specific selection scan.

X-axis refers to the number of genes, whereas Y-axis refers to individual GO terms. Colours indicate the adjusted *P*-values.

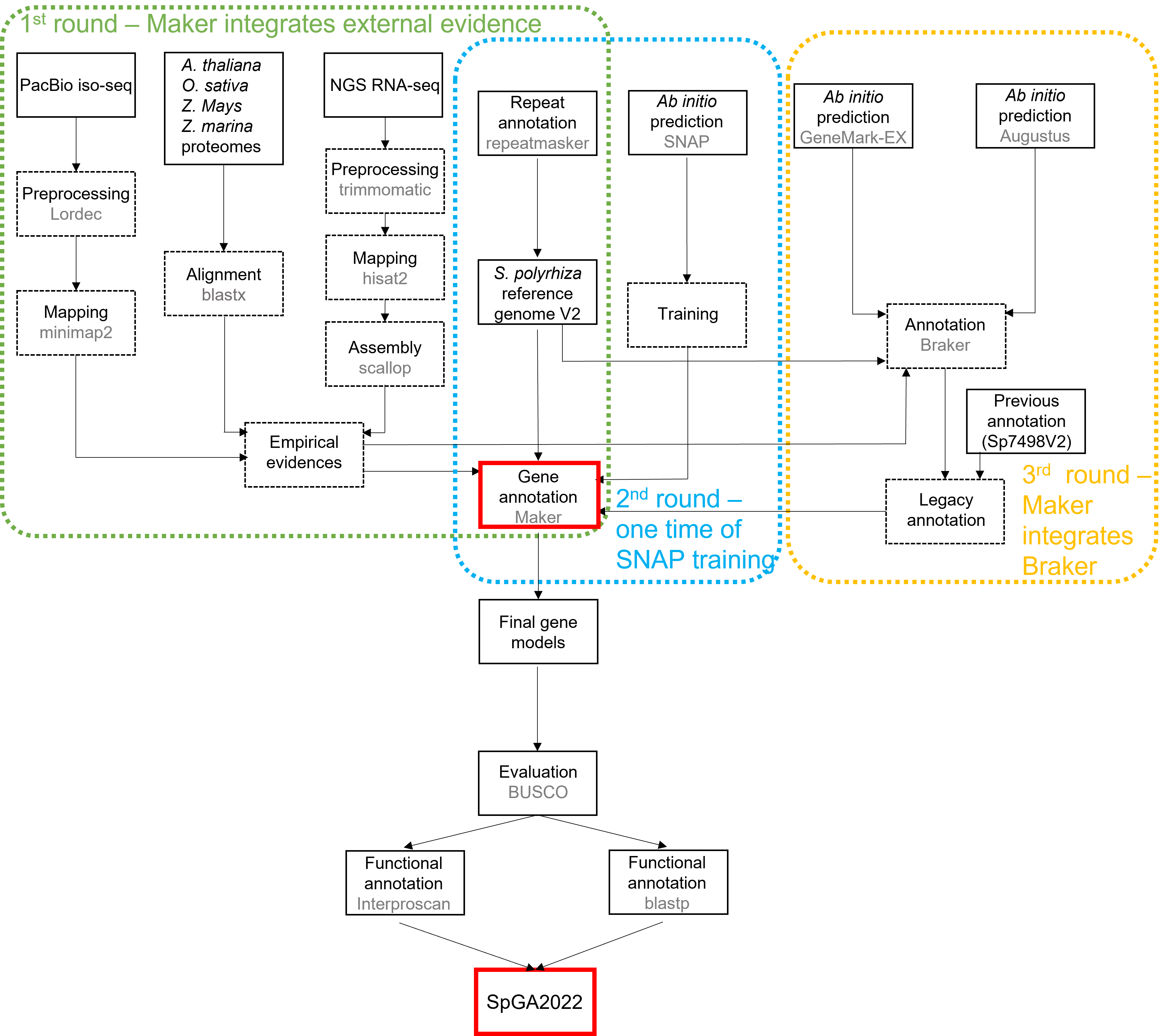

Supplementary Fig. 16. The schematic illustration of the genome annotation pipeline.

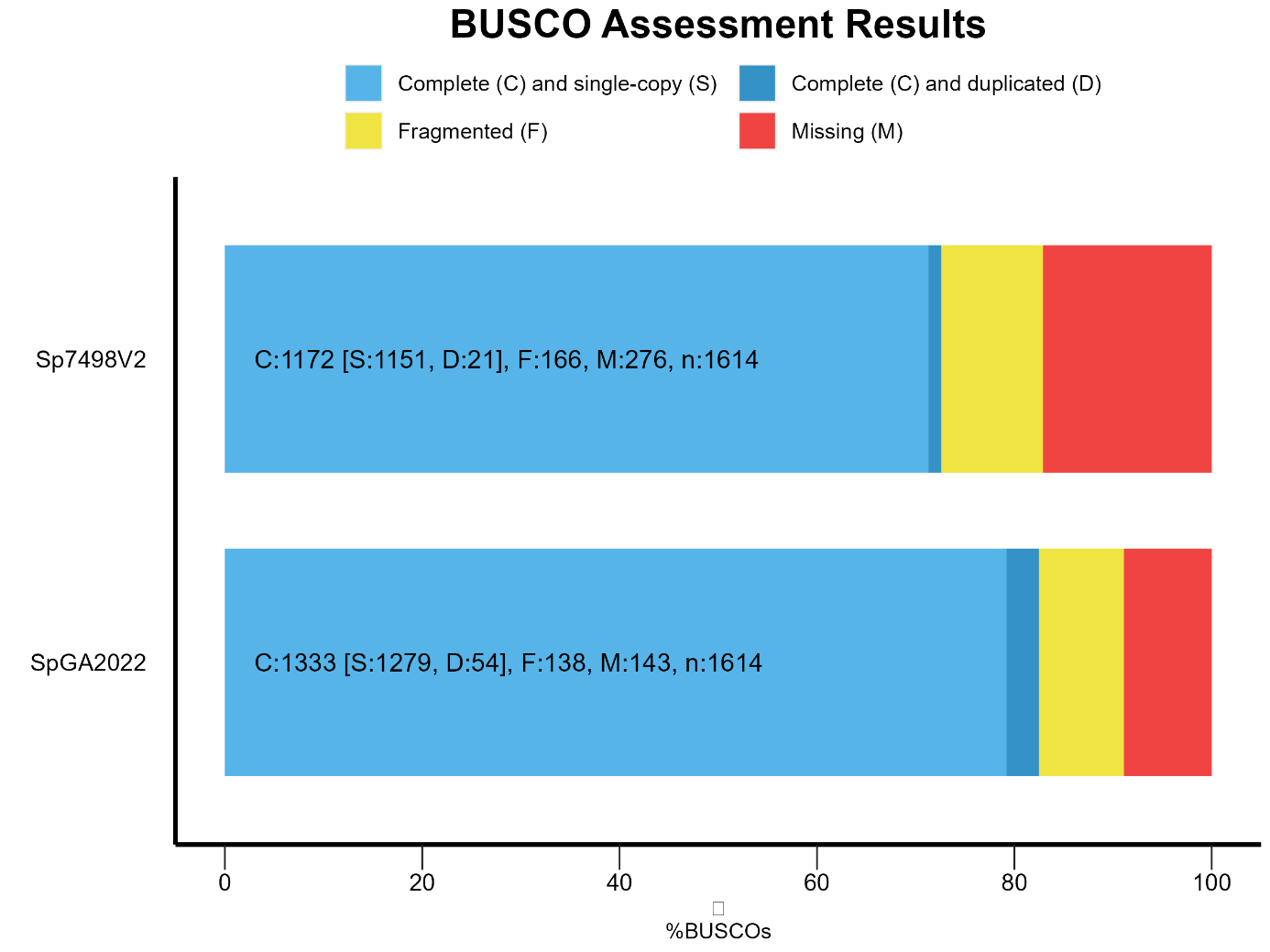

Supplementary Fig. 17. The BUSCO comparison between SpGA2022 and Sp7498V2.

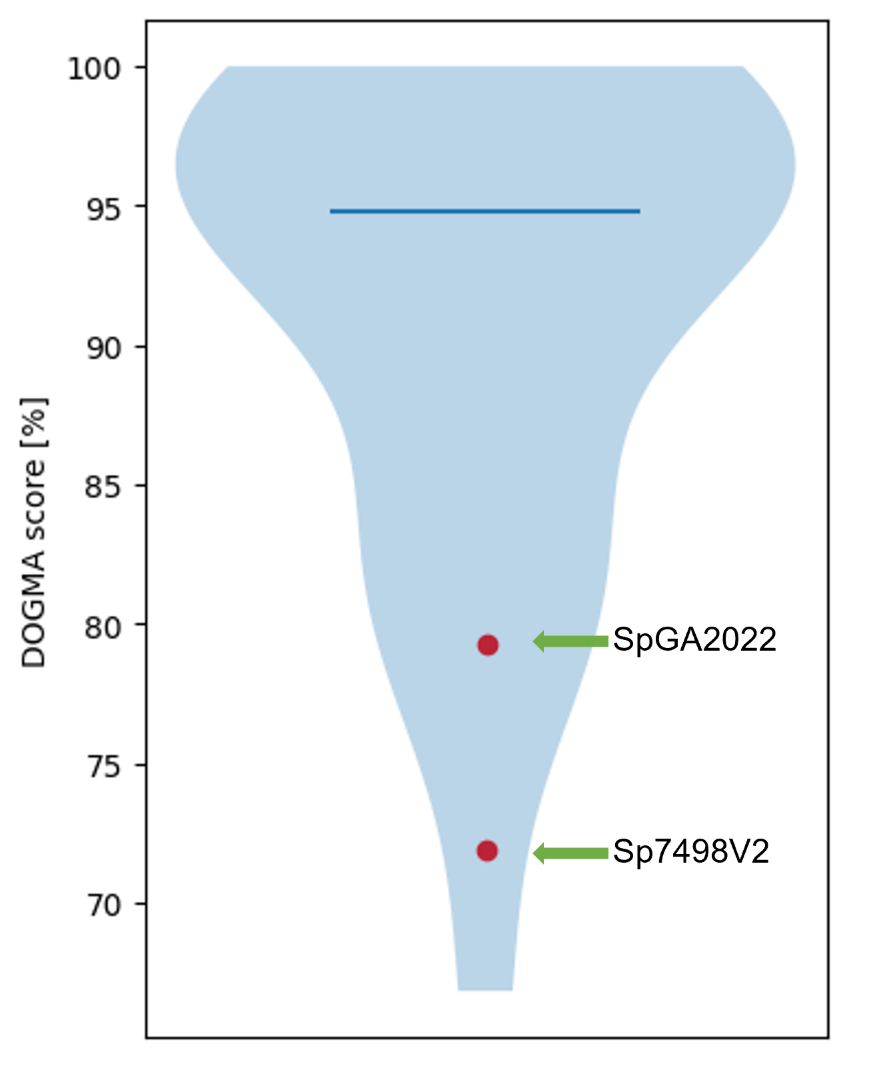

Supplementary Fig. 18. The DOGMA comparison between SpGA2022 and Sp7498V2.

The DOGMA scores of two annotations were shown in red points. The distribution of the other 23 monocot proteomes is shown in the blue area. The blue horizontal line indicates the median.

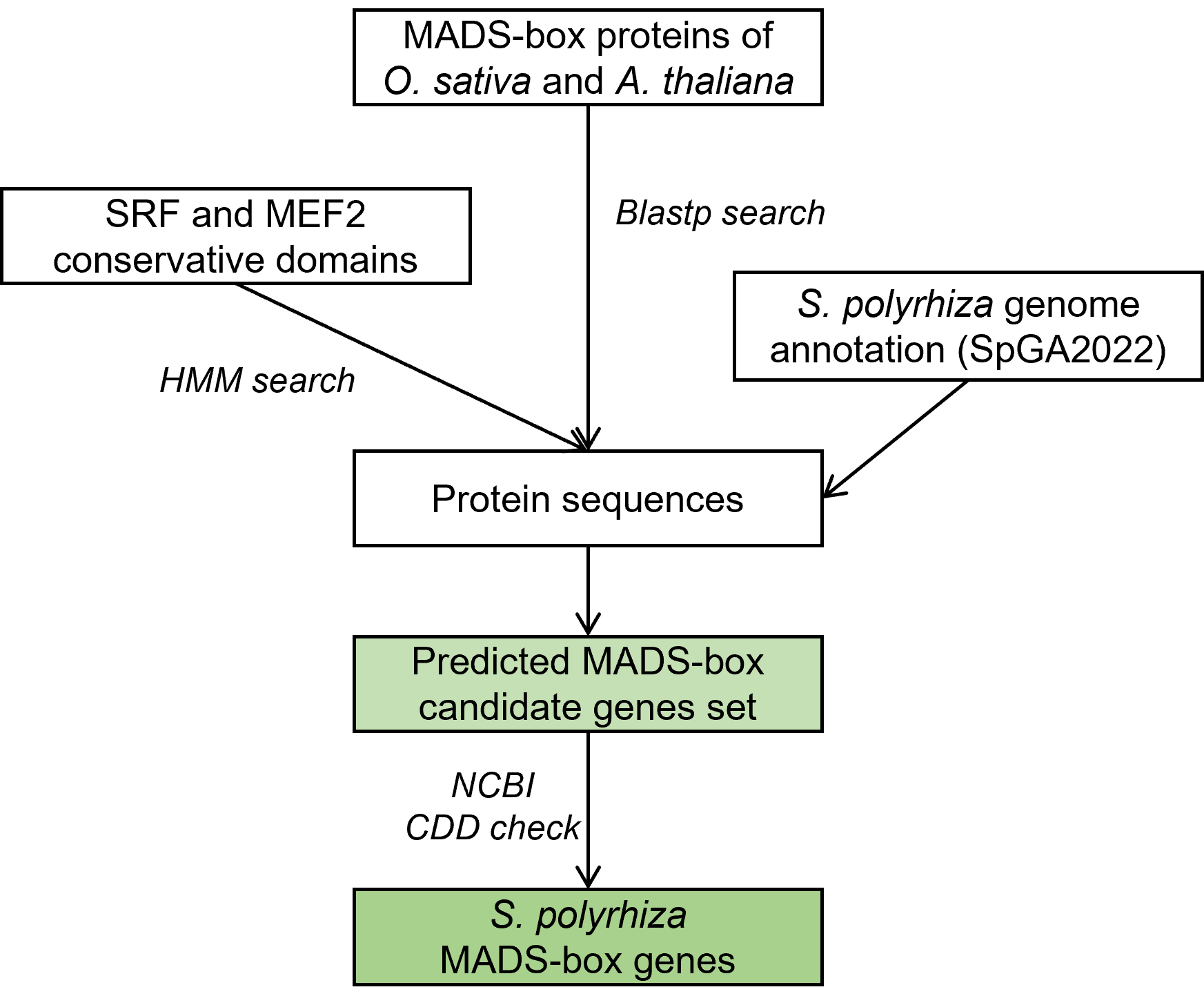

Supplementary Fig. 19. The *S. polyrhiza* MADS-box annotation pipeline.

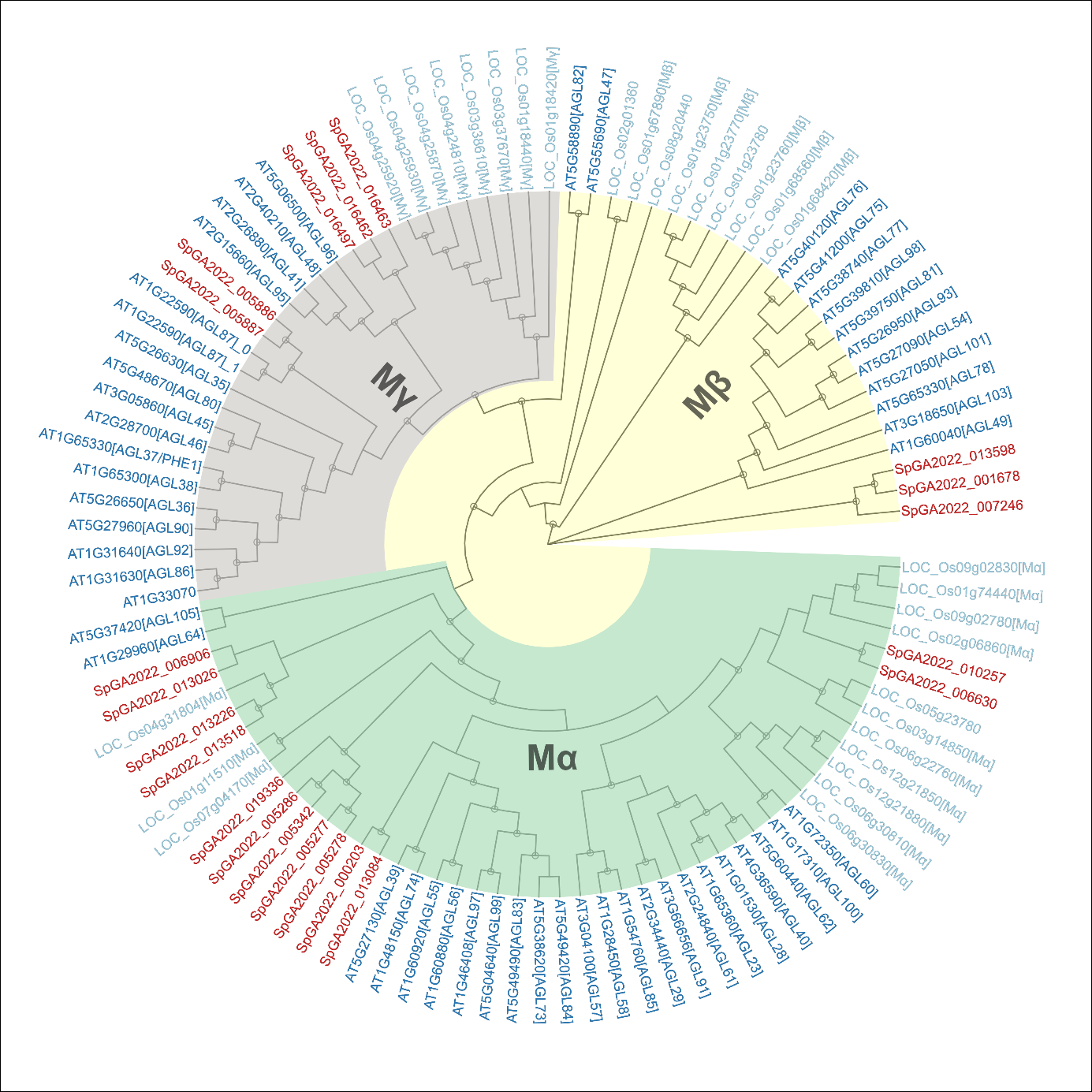

Supplementary Fig. 20. Phylogeny of Type I MADS-box transcription factors.

Phylogeny of Type I MADS-box transcription factor subfamily *in O. sativa*, *A. thaliana*, and *S. polyrhiza*. Three Type I MADS-box clades, Mα, Mβ, and Mγ, were highlighted using different colours. Tips are the gene names of *O. sativa* (starting with “LOC_Os” and indicated in light blue), *A. thaliana* (starting with “AT” and indicated in dark blue), and the *S. polyrhiza*’s updated gene annotation (SpGA2022, indicated in red). Genes from *O. sativa* and *A. thaliana* were partially annotated (information in square brackets) by previously classified clades based on public databases or publications (see methods). Small circles on the internal nodes suggest bootstrapping support higher than 0.75 (max of 1.00).

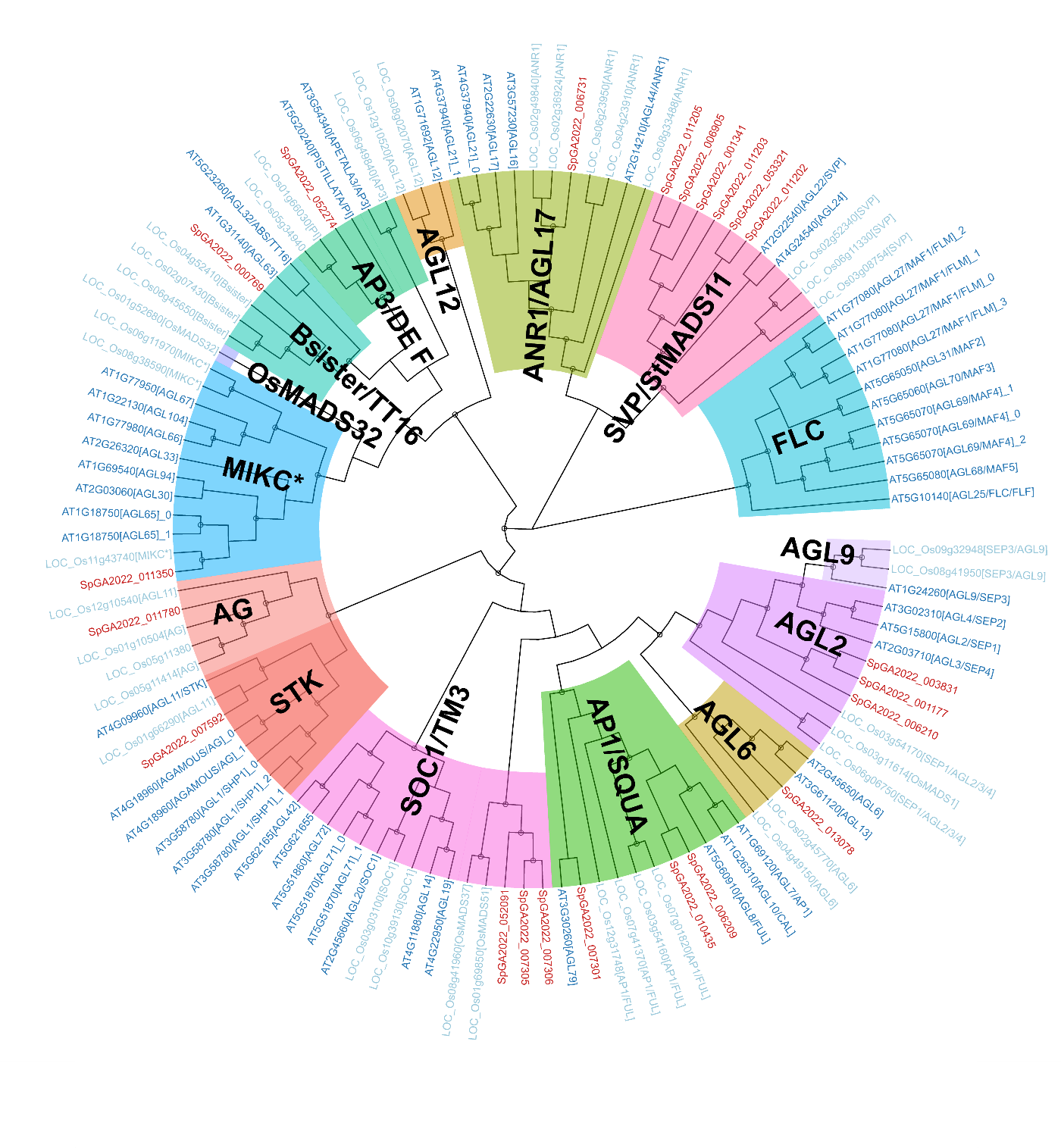

Supplementary Fig. 21. Phylogeny of Type II MADS-box transcription factors.

The phylogenetic relationship of 117 Type II MADS-box genes from three species was shown. The 15 subfamilies are marked in different colours.

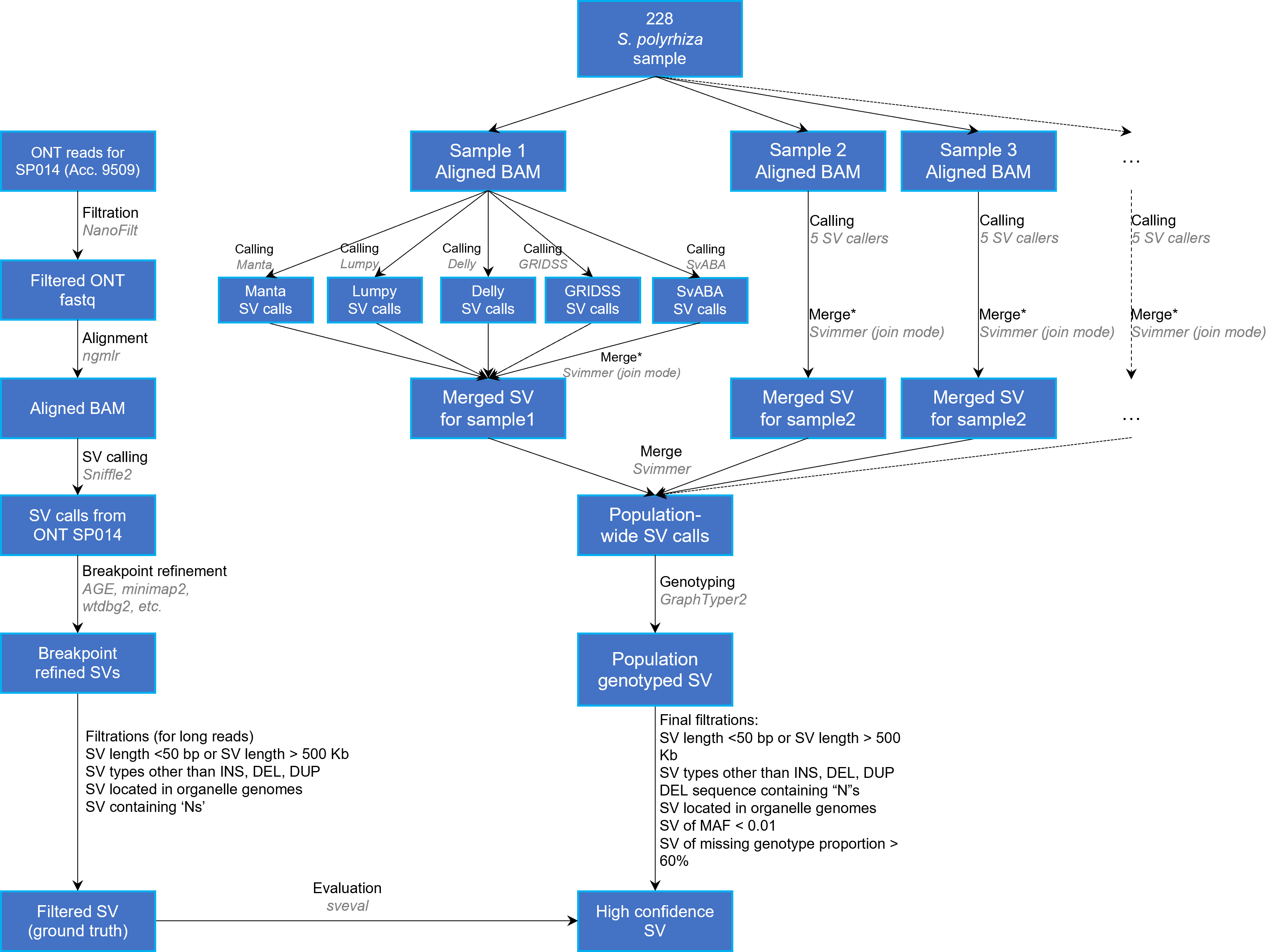

Supplementary Fig. 22. Structural variation calling pipeline.

Blue boxes indicate datasets or objects. Texts next to the arrows describe the processing steps. Tools used are shown in grey italic. *: merging SV calls from five different callers using Svimmer “join mode” means joining SV calls from Lumpy, Delly, GRIDSS, and SvABA with Manta.

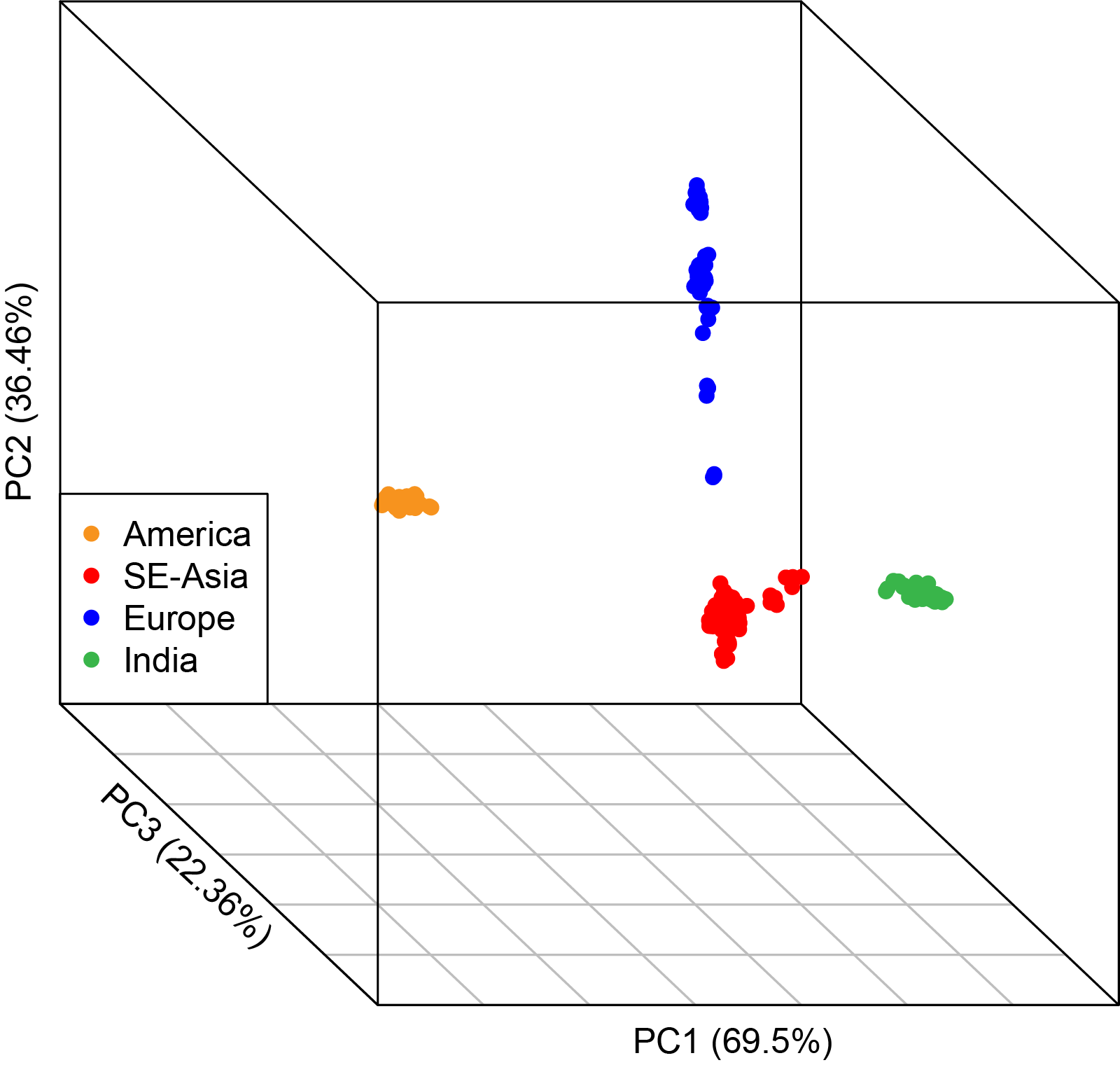

Supplementary Fig. 23. Principal component analysis based on SVs.

Three-dimensional plot showing the first three principal components. Small circular dots represent samples from the *S. polyrhiza* population. Different colours indicate the four populations that have been classified according to the population structure analysis.

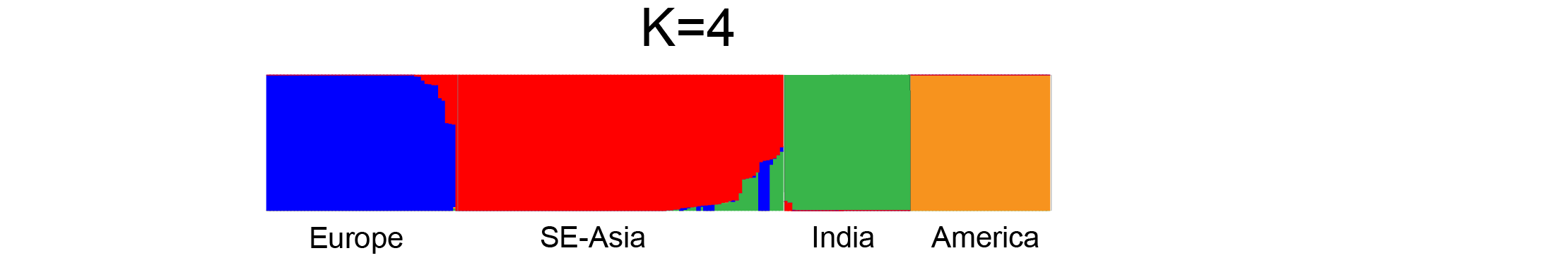

Supplementary Fig. 24. Population structure analyses based on SVs.

The plot shows the population structure analysis based on the SV data. The number of stratified groups was estimated as K=4 using fastStructure.

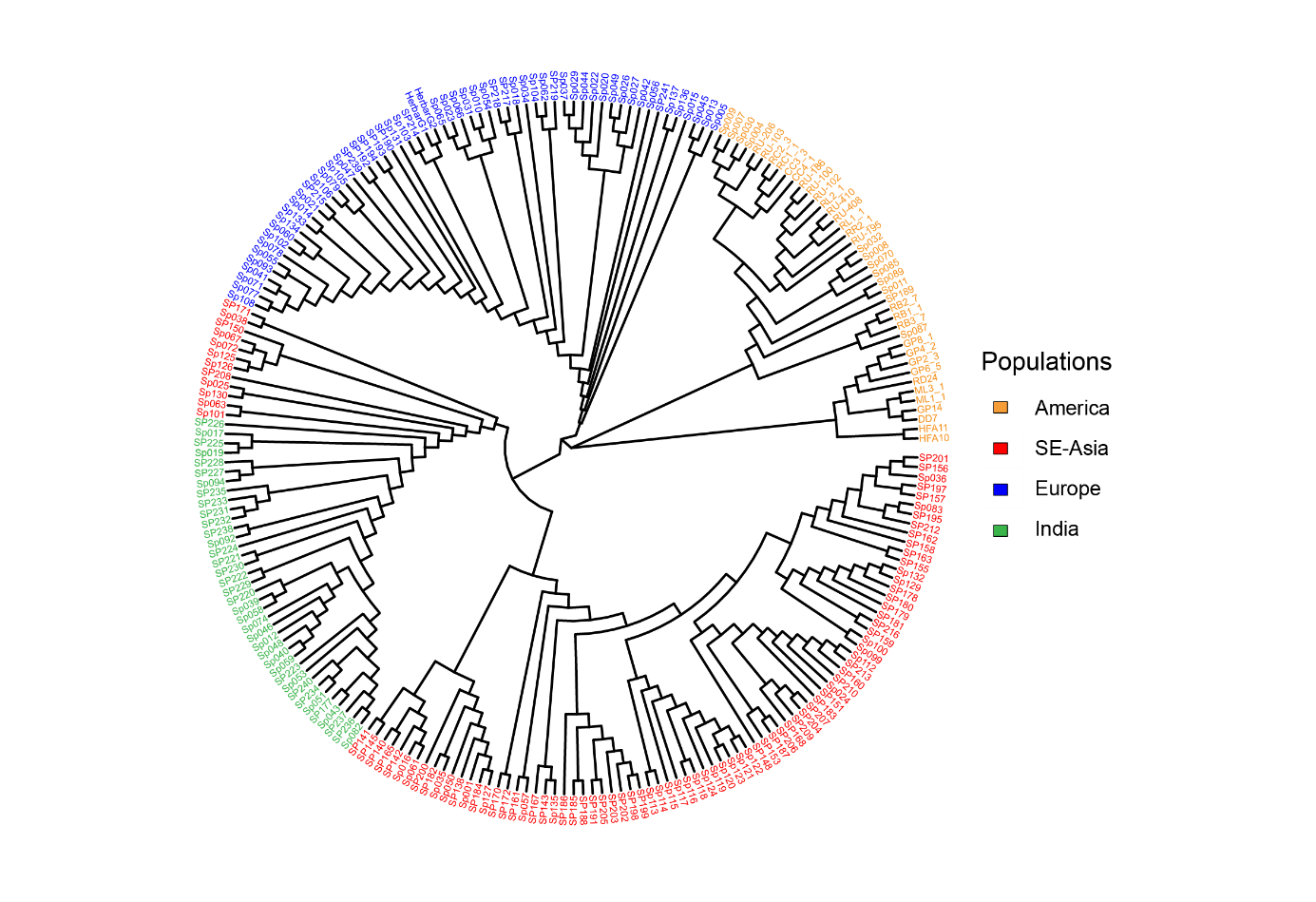

Supplementary Fig. 25. Neighbour-Joining phylogenetic tree based on SVs.

An unrooted neighbour-joining tree was reconstructed based on the p-distance calculated using population-level genotyped SVs. Tip names indicate sample names, while different colours represent the different populations to which the sample has been classified based on the population structure analysis: Asian (red), European (blue), Indian (green), and American (yellow).

**4. Supplementary Tables**

Supplementary Table 1. Pairwise nucleotide diversity (π) among different populations.

| **Population** | **π** | **π_N_*** | **π_S_*** | **π_N_/π_S_** |
| --- | --- | --- | --- | --- |
| America | 0.00045 | 0.00019 | 0.00044 | 0.424 |
| SE-Asia | 0.00113 | 0.00039 | 0.00106 | 0.367 |
| Europe | 0.00071 | 0.00026 | 0.00069 | 0.376 |
| India | 0.00061 | 0.00023 | 0.00056 | 0.410 |
| Combined | 0.00140 | 0.00049 | 0.0013 | 0.375 |

***:** π_N_ and π_S_ refer to the pairwise nucleotide diversity at nonsynonymous and synonymous sites, respectively.

Supplementary Table 2. The annotation of SVs.

|  | Insertions | Duplications | Deletions |
| --- | --- | --- | --- |
| 5’_UTR_variant | 15 | 75 | 87 |
| upstream_gene_variant | 298 | 63 | 591 |
| coding_sequence_variant | 34 | 155 | 169 |
| intron_variant | 303 | 209 | 919 |
| inframe_deletion | - | - | 20 |
| start_lost | - | - | 2 |
| start_retained_variant | - | - | 2 |
| stop_lost | - | - | 54 |
| frameshift_variant | - | - | 37 |
| downstream_gene_variant | 323 | 79 | 579 |
| 3’_UTR_variant | 22 | 80 | 99 |
| feature_elongation | 371 | 123 | - |
| feature_truncation | - | - | 1098 |
| intergenic_variant | 196 | 49 | 604 |
| transcript_ablation | - | - | 27 |
| splice_polypyrimidine_tract_variant | 5 | 111 | 112 |
| transcript_amplification | - | 509 | - |

Supplementary Table 3. The association of two MADS-box genes impacted SVs and the genome-wide heterozygosity rate within each population of *S. polyrhiza*.

| **SV** | **Chr** | **Position** | **Population** | **Beta** | **SE** | **P value** |
| --- | --- | --- | --- | --- | --- | --- |
| AGL62 (INS) | ChrS02 | 6841131 | SE-Asia | -0.0015 | 0.0046 | 0.745 |
|  |  |  | India | -0.00038 | 0.005 | 0.938 |
|  |  |  | Europe | 0.047 | 0.009 | **3.57e-07** |
| SOC1 (DEL) | ChrS04 | 5045470 | India | -0.0098 | 0.0047 | **0.037** |

Supplementary Table 4. The proportion of methylated cytosines among 20 *S. polyrhiza* methylomes.

| **Population** | **Sample** | **mC proportion (%)^#^** | | | |
| --- | --- | --- | --- | --- | --- |
|  |  | **C** | **CpG** | **CHG** | **CHH** |
| America | SP011 | 2.00 | 9.37 | 3.48 | 0.12 |
|  | SP032 | 1.88 | 9.19 | 3.07 | 0.10 |
|  | SP004 | 1.57 | 7.68 | 2.08 | 0.05 |
|  | SP085 | 1.33 | 6.43 | 2.01 | 0.07 |
|  | SP087 | 1.40 | 6.73 | 1.94 | 0.08 |
| SE-Asia | SP115 | 1.54 | 7.09 | 2.46 | 0.13 |
|  | SP150 | 1.74 | 7.88 | 2.21 | 0.16 |
|  | SP165 | 2.15 | 10.72 | 3.90 | 0.19 |
|  | SP182 | 1.27 | 5.37 | 1.79 | 0.14 |
|  | SP072 | 1.41 | 6.59 | 1.98 | 0.11 |
| Europe | SP015 | 1.19 | 6.45 | 0.82 | 0.10 |
|  | SP023 | 1.31 | 6.15 | 1.85 | 0.10 |
|  | SP031 | 1.77 | 8.36 | 3.28 | 0.17 |
|  | SP047 | 1.38 | 6.50 | 1.86 | 0.10 |
|  | SP055 | 1.51 | 6.87 | 2.53 | 0.12 |
| India | SP227 | 1.42 | 7.00 | 1.52 | 0.12 |
|  | SP236 | 1.30 | 6.31 | 1.58 | 0.11 |
|  | SP039 | 1.37 | 6.95 | 1.48 | 0.10 |
|  | SP043 | 1.26 | 5.91 | 1.54 | 0.11 |
|  | SP059 | 1.29 | 6.11 | 1.72 | 0.10 |

#mC proportion: The fraction of methylated cytosines in all sequencing-covered cytosines.

Supplementary Table 5. The list of 20 methylomes.

| **Clonal_family_ID** | **Sample_ID** | **Population** |
| --- | --- | --- |
| 9 | SP032 | America |
| 3 | SP087 | America |
| 11 | SP011 | America |
| 59 | SP085 | America |
| 5 | SP004 | America |
| 7 | SP047 | Europe |
| 10 | SP031 | Europe |
| 7 | SP055 | Europe |
| 39 | SP023 | Europe |
| 35 | SP015 | Europe |
| 145 | SP227 | India |
| 16 | SP039 | India |
| 47 | SP043 | India |
| 17 | SP059 | India |
| 154 | SP236 | India |
| 95 | SP165 | SE-Asia |
| 19 | SP072 | SE-Asia |
| 105 | SP182 | SE-Asia |
| 67 | SP115 | SE-Asia |
| 83 | SP150 | SE-Asia |

Supplementary Table 6. Comparison between annotations of SpGA2022 and Sp7498V2.

|  | **SpGA2022** | **Sp7498V2** |
| --- | --- | --- |
| # of genes | 20,546 | 19,620 |
| # of genes with UTR on both sides | 7,937 | - |
| # of genes with at least one UTR | 9,500 | - |
| # of single exon gene | 2,191 | 3,806 |
| Total CDS length | 23.6 Mb | 21.7 Mb |
| Total exon length | 30.5 Mb | 21.7 Mb |
| mean exons per mRNA | 5.5 | 5.2 |
| mean gene length | 4,225 | 3,455 |
| mean CDS length | 1,110 | 1,107 |
| mean exon length | 261 | 212 |
| mean five_prime_UTR length | 380 | - |
| mean three_prime_UTR length | 414 | - |
| # of mRNA without a start codon | 191 | 1,222 |
| # of mRNA without a stop codon | 470 | 508 |
| # of mRNA without both start and stop codons | 63 | 2,668 |

Supplementary Table 7. MADS-box transcription factors identification in *S. polyrhiza*.

| **SUB-FAMILIES** | **GENE_ID** | **CLADES** |
| --- | --- | --- |
| TypeI | SpGA2022_006630 | Mα |
| TypeI | SpGA2022_010257 | Mα |
| TypeI | SpGA2022_000203 | Mα |
| TypeI | SpGA2022_013084 | Mα |
| TypeI | SpGA2022_005286 | Mα |
| TypeI | SpGA2022_005342 | Mα |
| TypeI | SpGA2022_005277 | Mα |
| TypeI | SpGA2022_005278 | Mα |
| TypeI | SpGA2022_019336 | Mα |
| TypeI | SpGA2022_013226 | Mα |
| TypeI | SpGA2022_013518 | Mα |
| TypeI | SpGA2022_006906 | Mα |
| TypeI | SpGA2022_013026 | Mα |
| TypeI | SpGA2022_001678 | Mβ |
| TypeI | SpGA2022_013598 | Mβ |
| TypeI | SpGA2022_007246 | Mβ |
| TypeI | SpGA2022_005886 | Mγ |
| TypeI | SpGA2022_005887 | Mγ |
| TypeI | SpGA2022_016462 | Mγ |
| TypeI | SpGA2022_016497 | Mγ |
| TypeI | SpGA2022_016463 | Mγ |
| TypeII | SpGA2022_052274 | AP3/DEF |
| TypeII | SpGA2022_000769 | Bsister/TT16 |
| TypeII | SpGA2022_006731 | ANR1/AGL17 |
| TypeII | SpGA2022_001341 | SVP/StMADS11 |
| TypeII | SpGA2022_011205 | SVP/StMADS11 |
| TypeII | SpGA2022_011203 | SVP/StMADS11 |
| TypeII | SpGA2022_053321 | SVP/StMADS11 |
| TypeII | SpGA2022_011202 | SVP/StMADS11 |
| TypeII | SpGA2022_006905 | SVP/StMADS11 |
| TypeII | SpGA2022_011350 | MIKC* |
| TypeII | SpGA2022_007592 | AG/STK |
| TypeII | SpGA2022_011780 | AG/STK |
| TypeII | SpGA2022_052091 | SOC1/TM3 |
| TypeII | SpGA2022_007306 | SOC1/TM3 |
| TypeII | SpGA2022_007305 | SOC1/TM3 |
| TypeII | SpGA2022_010435 | AP1/SQUA |
| TypeII | SpGA2022_006209 | AP1/SQUA |
| TypeII | SpGA2022_007301 | AP1/SQUA |
| TypeII | SpGA2022_013078 | AGL6 |
| TypeII | SpGA2022_001177 | SEP/AGL2 |
| TypeII | SpGA2022_003831 | SEP/AGL2 |
| TypeII | SpGA2022_006210 | SEP/AGL2 |

Supplementary Table 8. List of MADS-box genes that are likely affected by structural variations.

| **Chr** | **Position** | **SV type** | **Size** | **Alternate Allele Frequency** | | | | **Affected genes** | **Annotations** |
| --- | --- | --- | --- | --- | --- | --- | --- | --- | --- |
|  |  |  |  | **America** | **SE-Asia** | **India** | **Europe** |  |  |
| 02 | 6832942 | DUP | 12832 | 0 | 0.083 | 0 | 0.291 | SpGA2022_005277^†^  SpGA2022_005278 | *AGL62* (Mα)  *AGL62* (Mα) |
| 02 | 6841131 | INS | 84 | 0 | 0.875 | 0.473 | 0.250^*^ | SpGA2022_005278 | *AGL62* (Mα) |
| 04 | 5003744 | DUP | 28875 | 0 | 0.097 | 0 | 0 | SpGA2022_052091^†^  SpGA2022_007305^†^ | *SOC1*  *SOC1* |
| 04 | 5045470 | DEL | 69 | 0 | 0 | 0.730 | 0 | SpGA2022_007306^#^ | *SOC1* |
| 07 | 8043832 | INS | 162 | 0.769 | 1 | 1 | 1 | SpGA2022_011350^#^ | *AGL65* (MIKC*) |
| 09 | 6604420 | DEL | 531 | 0 | 0.315 | 0 | 0.328 | SpGA2022_013518 | *AGL64* (Mα) |

^*^This insertion variation has missing genotyping higher than 60% in the European populations. For the rest of the SVs shown in the table, the overall missingness is lower than 13%.

^†^Those genes overlapped with SVs in the genic region and the upstream 2 kb region.

^#^Those genes overlapped with SVs only in the upstream 2 kb regions.

Supplementary Table 9. Inferred demographic parameters.

| Parameter | Prior | Posterior | 95% Confidence intervals |
| --- | --- | --- | --- |
| *N*e_Anc_ | *U*(1x10^4^,4x10^6^) | 44875 ind. | (41853,48497) |
| *N*e_AME_ | *U*(0.3,50)**N*e_Anc_ | 8.716**N*e_Anc_ (~400000 ind.) | (0.3,24.83) |
| *N*e_ASIA_ | *U*(0.3,50)**N*e_Anc_ | 14.652**N*e_Anc_ (~660000 ind.) | (0.3,45.4) |
| *N*e_EUR_ | *U*(0.3,3)**N*e_Anc_ | 1.657**N*e_Anc_ (~75000 ind.) | (0.83,2.43) |
| *N*e_IND_ | *U*(0.3,3)**N*e_Anc_ | 3.000**N*e_Anc_ (~135000 ind.) | (0.3,3) |
| *T*_AME_ | *U*(*T*_IND_,30)**N*e_Anc_ | 22.021**N*e_Anc_ (~1x10^6^ gen. ago) | (0.0001,27.43) |
| *T*_ASIA_ | *U*(*T*_IND_,30)**N*e_Anc_ | 22.454**N*e_Anc_ (~1x10^6^ gen. ago) | (18.39,23.26) |
| *T*_EUR_ | *U*(0.0001,1)**N*e_Anc_ | 0.266**N*e_Anc_ (~12000 gen. ago) | (0.13,0.60) |
| *T*_IND_ | *U*(*T*_EUR_,30)**N*e_Anc_ | 1.120**N*e_Anc_ (~51000 gen. ago) | (0.0001,18.19) |
| *M*_ASIA_ | *U*(10^-13^,10^-1^) | 4*N*e_Anc_*m* = 7.4302x10^-9^ | (10^-11.9^,10^-6.1^) |
| *M*_EUR_ | *U*(10^-13^,10^-1^) | 4*N*e_Anc_*m* = 1.3932x10^-5^ | (10^-9.41^,10^-3.28^) |
| *M*_IND_ | *U*(10^-13^,10^-1^) | 4*N*e_Anc_*m* = 5.1286x10^-13^ | (10^-13^,10^-10.20^) |

Supplementary Table 10. Intergenic loci selected for demographic inference.

| **Chromosome** | **From** | **To** | **Size (bp)** |
| --- | --- | --- | --- |
| ChrS01 | 10341926 | 10370644 | 28718 |
| ChrS01 | 6201042 | 6245822 | 44780 |
| ChrS02 | 4580335 | 4614088 | 33753 |
| ChrS02 | 7338329 | 7371076 | 32747 |
| ChrS03 | 2135139 | 2171492 | 36353 |
| ChrS03 | 3324230 | 3353948 | 29718 |
| ChrS04 | 2398564 | 2428164 | 29600 |
| ChrS04 | 5843951 | 5877409 | 33458 |
| ChrS04 | 8231251 | 8276638 | 45387 |
| ChrS05 | 5717537 | 5749587 | 32050 |
| ChrS05 | 5837287 | 5867849 | 30562 |
| ChrS06 | 2104286 | 2154345 | 50059 |
| ChrS06 | 2203817 | 2245642 | 41825 |
| ChrS07 | 4834724 | 4874602 | 39878 |
| ChrS09 | 1416036 | 1445139 | 29103 |
| ChrS09 | 3633841 | 3670635 | 36794 |
| ChrS09 | 3947117 | 3977500 | 30383 |
| ChrS10 | 6686084 | 6733993 | 47909 |
| ChrS11 | 2990169 | 3034837 | 44668 |
| ChrS11 | 5005366 | 5045753 | 40387 |
| ChrS12 | 2218087 | 2244224 | 26137 |
| ChrS12 | 457799 | 491403 | 33604 |
| ChrS13 | 1568966 | 1600801 | 31835 |
| ChrS13 | 5071361 | 5102204 | 30843 |
| ChrS14 | 1987237 | 2018224 | 30987 |
| ChrS14 | 340148 | 375438 | 35290 |
| ChrS14 | 4545337 | 4578899 | 33562 |
| ChrS15 | 1710444 | 1752107 | 41663 |
| ChrS15 | 1753520 | 1781319 | 27799 |
| ChrS15 | 3867569 | 3897413 | 29844 |
| ChrS16 | 1261209 | 1301988 | 40779 |
| ChrS16 | 3523583 | 3556154 | 32571 |
| ChrS16 | 4504899 | 4543897 | 38998 |
| ChrS17 | 2732045 | 2771792 | 39747 |
| ChrS18 | 2486433 | 2511566 | 25133 |
| ChrS18 | 664864 | 689461 | 24597 |
| ChrS19 | 2569311 | 2610161 | 40850 |
| ChrS20 | 1129392 | 1151473 | 22081 |
| ChrS20 | 715644 | 745921 | 30277 |
