## Supplementary material for "Population genomics and epigenomics provide insights into the evolution of facultative asexuality in plants": Description of Additional Supplementary Files.docx

**Supplementary Data 1:** The file describes detailed information on all 228 *S. polyrhiza* genotypes, including their sequencing statistics, sampling regions, and Accession number in NCBI.

**Supplementary Data 2:** The table contains the annotation of all SNPs. Each row records one annotated gene. The first column is the gene ID, and the following columns are the number of SNPs annotated as different categories.

**Supplementary Data 3:** The table contains the grouping of clonal families of all 228 *S. polyrhiza* genotypes.

**Supplementary Data 4:** The table contains the list of genes detected under positive selection at the species-wide level according to RAiSD, SweeD and LASSI.

**Supplementary Data 5:** This table contains the list of genes (rows) belonging to the top 1% CLR values of 3P-CLR and the population branches where they are under selection (columns). Fields with “1” indicate that a signature of selection was found, and fields with “0” indicate no signature of selection was found.

**Supplementary Data 6:** The file contains detailed information for the NCBI CDD check of annotated MADS-box TFs. The first sheet records all 43 MADS-box TFs that passed the CDD check, while the second sheet records other candidates that failed the CDD check.
